# Development of a high-throughput pipeline to characterize microglia morphological states at a single-cell resolution

**DOI:** 10.1101/2023.11.03.565581

**Authors:** Jennifer Kim, Paul Pavlidis, Annie Vogel Ciernia

## Abstract

As rapid responders to their environments, microglia engage in functions that are mirrored by their cellular morphology. Microglia are classically thought to exhibit a ramified morphology under homeostatic conditions which switches to an ameboid form during inflammatory conditions. However, microglia display a wide spectrum of morphologies outside of this dichotomy, including rod-like, ramified, ameboid, and hypertrophic states, which have been observed across brain regions, neurodevelopmental timepoints, and various pathological contexts. We used dimensionality reduction and clustering approaches to consider contributions of multiple morphology measures together to define a spectrum of microglial morphological states. Using ImageJ tools, we first developed a semi-automated approach to characterize 27 morphology features from hundreds to thousands of individual microglial cells in a brain subregion-specific manner. Within this pool of morphology measures, we defined distinct sets of highly correlated features that describe different aspects of morphology, including branch length, branching complexity, territory span, and cell circularity. When considered together, these sets of features drove different morphological clusters. Furthermore, our analysis toolset captured morphological states similarly and robustly when applied to independent datasets and using different immunofluorescent markers for microglia. We have compiled our morphology analysis pipeline into an accessible, easy to use, and fully open-source ImageJ macro and R package that the neuroscience community can expand upon and directly apply to their own analyses. Outcomes from this work will supply the field with new tools to systematically evaluate the heterogeneity of microglia morphological states across various experimental models and research questions.

## 1 Introduction

Clinical and postmortem studies support a role for altered microglial function as a critical component of multiple brain disorders, including Schizophrenia, Autism Spectrum Disorder, and Alzheimer’s Disease. (Hansen et al., 2018; Suzuki et al., 2013; Tetreault et al., 2012; Zhuo et al., 2023) In addition to their resident immune functions, microglia play critical roles to establish and maintain normal brain function including the regulation of neuronal cell number (Cunningham et al., 2013), shaping of brain circuitry (Bialas & Stevens, 2013; Schwarz et al., 2012), and fine-tuning of neuronal connections (Schafer et al., 2012; Wang et al., 2020; Zhan et al., 2014), processes that have all been shown to be fundamentally disrupted in many brain disorders. (Hammond et al., 2018; Lenz & Nelson, 2018) Microglia directly communicate with other cell types and modulate their function by releasing and responding to various molecular substrates in the brain environment including cytokines, chemokines, and neurotransmitters (Salvador et al., 2021). Thus, dysregulated microglial responses can disrupt the homeostatic brain environment and normal communication across cell types, ultimately altering brain function and behavior.

As rapid responders to their local environments, microglia exhibit a dynamic range of phenotypes defined by multiple parameters including transcriptomic signatures (Hammond et al., 2019; Li et al., 2019), epigenomic regulation (Ciernia et al., 2018; Meleady et al., 2023), and **changes in cellular morphology**. Microglia are classically thought to engage in immune functions that are mirrored in morphology, where homeostatic microglia exhibit a ramified morphology that switches to an ameboid, unramified form in response to inflammatory signals in their environment. This morphological switch is thought to allow for increased mobility to sites of infection or injury, efficient phagocytosis, and release of cytokines into the microenvironment, all functions that are characteristic of a “pro-inflammatory” state. However, a mechanistic link between reduced branching and increased inflammatory function has only been shown recently, where increased P2RY12 potentiation of the THIK-1 channel caused decreased microglial ramifications and increased activity of the Il1b inflammasome in response to tissue damage (Madry et al., 2018, p. 1; Paolicelli et al., 2022). Cdk1-mediated microtubule remodeling has also recently been shown to be required for efficient cytokine trafficking and release and transformation of microglia from ramified to ameboid forms after LPS exposure *in vitro* and *in situ*. (Adrian et al., 2023) In contrast, findings from other studies display a reverse relationship, where ramified microglia have been shown to phagocytose synapses during adult neurogenesis (Paolicelli et al., 2022; Sierra et al., 2010) and ameboid microglia show reduced phagocytic capabilities in epilepsy (Abiega et al., 2016; Paolicelli et al., 2022). The relationship between form and function is clearly not as dichotomous as historically thought, and there has been an increasing effort in the field to move away from dualistic classification of microglial function and towards a clearer understanding and appreciation of the heterogenous states of microglia (Dubbelaar et al., 2018; Paolicelli et al., 2022).

Microglia morphology has been shown to be highly context and signal-dependent, displaying various forms ranging from ameboid-like under inflammatory states to hyper-ramified in mouse models of stress-induced depression, accelerated aging, and Alzheimer’s disease (Beynon & Walker, 2012; Hellwig et al., 2016; Madry et al., 2018; Raj et al., 2014). Subsets of microglial populations have also been shown to display rod-like morphologies characterized by long, thin processes protruding from oval-shaped somas that retract neuron-adjacent planar processes in response to diffuse brain injury (Taylor et al., 2014). Furthermore, microglia are highly motile and never truly quiescent even in healthy conditions, constantly extending out protrusions to scan their environments for pathogens and other harmful signals, as revealed by *in vivo* two-photon imaging studies of the mouse brain (Bernier et al., 2020; Davalos et al., 2005; Nimmerjahn et al., 2005). Different morphological forms have been observed to be conserved across species (Paolicelli et al., 2022) and spatially distributed across brain regions, neurodevelopmental timepoints, and various pathological contexts (Savage et al., 2019) . Microglial morphological forms including ramified, rod-like, reactive or hypertrophic (rounder cell body with fewer and shorter processes) (Paolicelli et al., 2022), and ameboid (less than two unramified processes) (Paolicelli et al., 2022) (Fig. 1B) have been commonly observed in both humans and mice, and microglia have been shown to display similar dendritic morphology across species. (Geirsdottir et al., 2019; Paolicelli et al., 2022)

**Figure 1:**
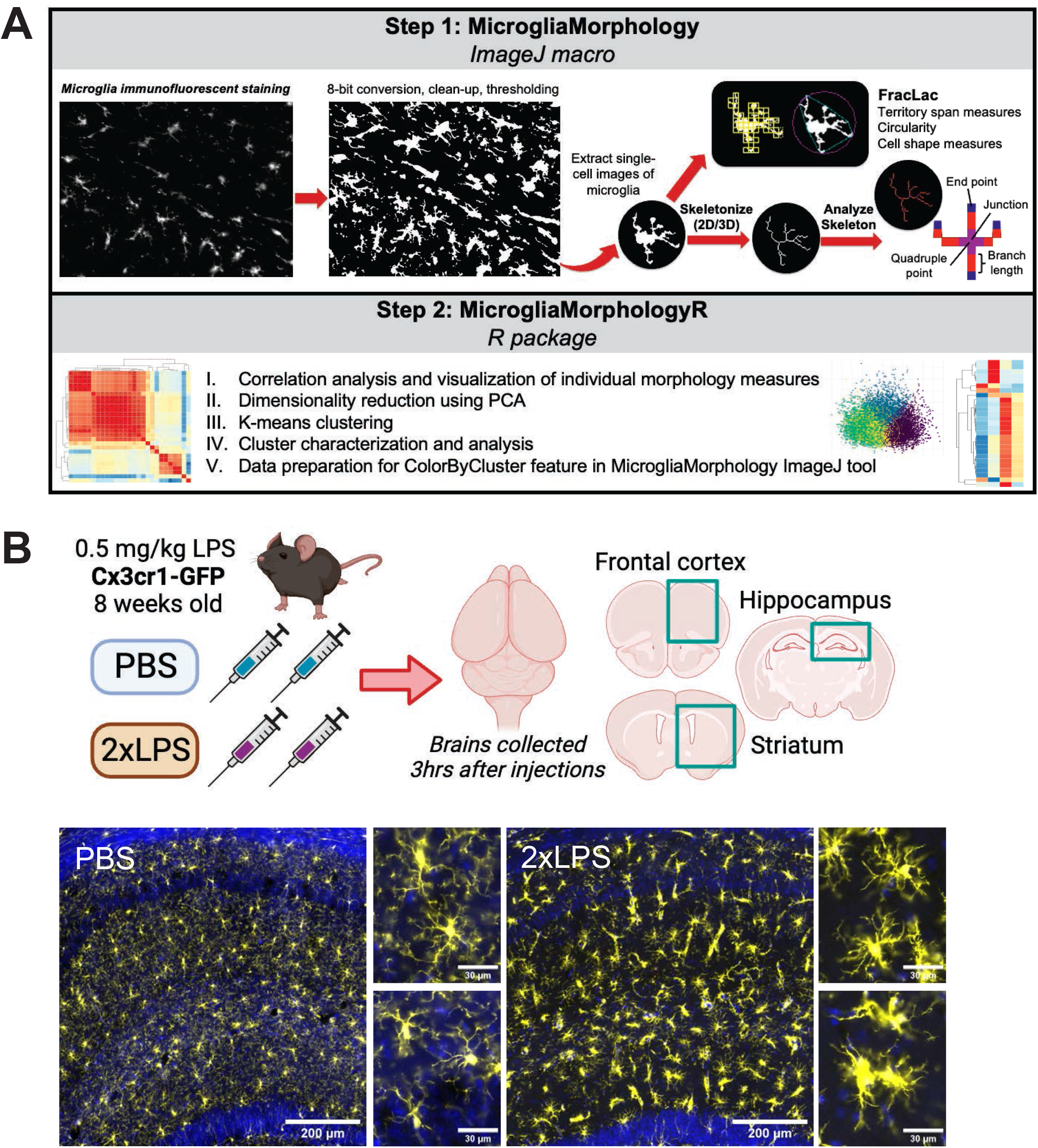
Study Overview. (A) Outline of steps involved in MicrogliaMorphology and MicrogliaMorphologyR. (B) Experimental mouse model used for dataset described throughout paper. Example images from dorsal hippocampus with individual microglia insets for each treatment condition. Microglia (Iba1) in yellow and DAPI nuclear stain in blue. Full size images scale bar 200um, insets 30um.

Microglia morphology can be explained using various measures of branch length, branching complexity, territory span, and cell circularity, which together comprise distinct sets of morphology measures that are changing in conjunction with each other. Nevertheless, studies often selectively report changes in individual features such as the number of branches or cell area alone, ultimately depicting an incomplete or biased representation of changes in microglia morphology that fail to capture true morphological states. Therefore, an analysis approach that considers contributions of all feature measures together to explain various axes of morphology is necessary to gain a better understanding of a microglia’s actual morphological state and relationship to cellular function. A plethora of tools exist to analyze microglia morphology (Clarke et al., 2021; Salamanca et al., 2019; York et al., 2018; Young & Morrison, 2018), but often involve significant effort to extract meaningful information at a larger scale, as many approaches that allow for the analysis of morphological information at a cellular resolution involve manually choosing and segmenting individual cells within an image. This introduces a potential bias for which cells are selected and vastly limits the feasible sample size for analysis. While there has been considerable progress in the development of toolsets which automate these time-consuming steps, there has been less development, transparency about, and availability of methods to analyze the resulting morphological measures (Reddaway et al., 2023). The underlying code and datasets for the majority of published toolsets are not openly available nor well-documented, further limiting the uptake and progression of the most up-to-date toolsets by the larger research community. (Reddaway et al., 2023)

Here, we describe an **accessible and open-source** morphology analysis toolset: MicrogliaMorphology (ImageJ tool) and MicrogliaMorphologyR (R package), which supplies the field with new tools to systematically evaluate the heterogeneity of microglia morphological states by considering 27 different measures of microglia morphology. To demonstrate use cases for our toolset, we characterized and analyzed microglia morphology in an experimental model of repeated immune stimulation by peripheral lipopolysaccharide (LPS) administration, a commonly used model (H. Jung et al., 2023; Wendeln et al., 2018) which induced population shifts in the four major classes of microglia morphology in our dataset: ramified, hypertrophic, ameboid, and rod-like. Application of MicrogliaMorphology and MicrogliaMorphologyR by the scientific community will yield novel insights into microglia morphology differences in the brain at a single-cell resolution and in a spatially-resolved manner across various experimental models and research questions. Additionally, the toolset is not limited to microglia morphology alone, but can also be applied in the same way to characterize morphology of other cell types.

## 2 Materials and Methods

### In vivo experiments

All experiments were conducted in accordance with the Canadian Council on Animal Care guidelines, with approval from the University of British Columbia’s Animal Care Committee. Mice were housed in groups of two to four on a regular 12-hr light/12-hr dark cycle and all experiments were performed in regular light during the mouse’s regular light cycle. CX3C motif chemokine receptor 1 (Cx3cr1) is a commonly used microglial marker that is expressed on microglia and other immune cells, and cells under control of the endogenous Cx3cr1 locus express GFP (IMSR_JAX:005582). (S. Jung et al., 2000) 8-week old male and female Cx3cr1-GFP mice bred on a C57BL/6J background (n=2 females and 1 male/condition) were intraperitoneally injected with 0.5 mg/kg lipopolysaccharide (LPS; Lipopolysaccharides from *E. coli* O55:B5, Sigma-Aldrich L5418) or vehicle solution (PBS; Phosphate buffered saline, Fisher BioReagents BP3991) once every 24 hours for 2 days.

**1xLPS experiments (**Supplementary Figure 3): Mice were housed in groups of two to four on a reversed 12-hr light/12-hr dark cycle. Because mice are nocturnal animals, all experiments were performed in red light during the mouse’s dark cycle when they are most active. 8-week old male and female C57BL/6J mice (n=2/sex/condition) were intraperitoneally injected once with 1.0 mg/kg lipopolysaccharide (LPS; Lipopolysaccharides from *E. coli* O55:B5, Sigma-Aldrich L5418) or vehicle solution 1xPBS (Phosphate buffered saline, Fisher BioReagents BP3991).

### Tissue collection

3 hours (2xLPS experiments) or 24 hours (1xLPS experiments) after the final injection, mice were quickly anesthetized with isofluorane and transcardially perfused with 15ml of 1xPBS before brains were extracted for downstream immunohistochemistry experiments. Extracted brains were immersion-fixed in 4% paraformaldehyde for 48 hours before cryoprotecting in 30% sucrose for 48 hours prior to cryosectioning. Cryoprotected brains were then sectioned at 30um on the cryostat, collected in 1xPBS for long-term storage, and processed for immunohistochemistry.

### Immunohistochemistry

30um brain sections were immunofluorescently stained for various markers of microglia: ionized calcium binding adaptor molecule 1 (Iba1) and/or purinergic receptor P2Y12 (P2ry12) to analyze microglial morphology. Brain sections in the 1xLPS experiments (Supp. Fig. 3) were stained with only Iba1 and brain sections in the 2xLPS experiments were stained with both Iba1 and P2ry12. Free-floating brain sections were washed 3 times for 5 minutes each in 1xPBS, permeabilized in 1xPBS + .5% Triton (Fisher BioReagents BP151-500) for 5 minutes, and incubated in blocking solution made of 1xPBS + .03% Triton + 1% Bovine Serum Albumin (BSA; Bio-techne Tocris 5217) for 1 hour. After the blocking steps, sections were incubated overnight at 4°C in primary antibody solution containing 2% Normal Donkey Serum (NDS; Jackson Immunoresearch Laboratories Inc. 017-000-121) + 1xPBS + .03% Triton + primary antibodies (chicken anti-Iba1: 1:1000, Synaptic Systems 234 009; rabbit anti-P2RY12: 1:500, Anaspec AS-55043A). After primary antibody incubation, sections were washed 3 times for 5 minutes each with 1xPBS + .03% Triton before incubating for 2 hours in secondary solution containing 2% NDS + DAPI (1:1000; Biolegend 422801) + secondary antibodies (Alexafluor 647 donkey anti-chicken: 1:500, Jackson Immunoresearch Laboratories Inc. 703-605-155; Alexafluor 568 donkey anti-rabbit: 1:500, Invitrogen A10042). Sections were washed 3 times for 5 minutes each with 1xPBS + .03% Triton before being transferred into 1xPBS for temporary storage before mounting. Sections were mounted onto microscope slides (Premium Superfrost Plus Microscrope Slides, VWR CA48311-703) and air dried before being coverslipped with mounting media (ProLong Glass Antifade Mountant, Invitrogen P36980; 24×60mm 1.5H High Performance Coverslips, Marienfield 0107242).

### Imaging

All mounted brain sections were imaged on the ZEISS Axioscan 7 microscope slide scanner at 20x magnification with a step-size of 0.5um using the z-stack acquisition parameters within the imaging software (ZEISS ZEN 3.7). During image acquisition, Extended Depth of Focus (EDF) images were created using maximum projection settings and saved as the outputs in the final .czi files. Maximum projection EDF images only compile the pixels of highest intensity at any given position in a z-stack to construct a new 2D image which retains the 3D information. Using ImageJ, we created .tiff images of each fluorescent channel from .czi files and selected and saved .tiffs of brain regions of interest (ROIs) to use as input for downstream morphological analysis in MicrogliaMorphology and MicrogliaMorphologyR. We focused our analyses on multiple brain regions including the hippocampus, frontal cortex, and striatum, as well as subregions within them. Images of coronal brain sections containing these regions were aligned to the Allen Brain Atlas (mouse.brain-map.org) (Pinskiy et al., 2015) using the ImageJ macro FASTMAP (Terstege et al., 2022).

### MicrogliaMorphology

MicrogliaMorphology is designed to be a user-friendly ImageJ macro that wraps around existing ImageJ plugins AnalyzeParticles, Skeletonize (2D/3D), and AnalyzeSkeleton, and is written using the ImageJ macro (IJM) language. (Schneider et al., 2012) All supporting code for MicrogliaMorphology is available on Github at https://github.com/ciernialab/MicrogliaMorphology and a detailed video tutorial which includes relevant troubleshooting steps is available on Youtube at https://www.youtube.com/watch?v=YhLCdlFLzk8. After MicrogliaMorphology is installed into the user’s ImageJ plugins folder as described in the Github repository, it will appear in the Plugins dropdown menu from the ImageJ toolbar, where it can be clicked on to begin the user prompts. In Step 1, users are be prompted to measure dataset-specific parameters, which are critical because every imaging dataset is prepared and acquired differently, and thus requires user input to determine what parameters most appropriately and accurately capture microglia morphology within individual datasets. Image thresholding parameters including the method and radius considered for auto local thresholding are important to determine the most appropriate thresholding method which captures full, single microglial cells without losing branching connectivity. Area ranges that accurately describe single microglial cells are important to exclude any artifacts of 2D representation in the EDFs such as cell particles and incomplete cells or multiple cells that are overlapping. Dataset-specific image thresholding and cell area parameters determined by these initial steps are then called to within the macro to inform downstream steps of MicrogliaMorphology. In the subsequent steps, the only user input involves following prompts to select input folders to call from and output folders to write to, with the option of batch-processing. All protocols, computation, and analysis described below have been written to be automated within MicrogliaMorphology, unless otherwise specified.

Step 2 after determining dataset-specific parameters is thresholding input .tiff fluorescent images. MicrogliaMorphology cleans up and thresholds input images according to standard protocol (Young & Morrison, 2018): images are binarized and converted to grayscale before the brightness and contrast are enhanced, unsharp mask filter applied to clarify existing detail, despeckle function applied to remove noise, and auto local or auto thresholding applied. Then, a second despeckle step is applied before dilation and connection steps are applied to connect branches, after which outliers are removed. The final images are then used as input for Step 3, which uses AnalyzeParticles and ROI manager functions to create and save new images of single-cells which pass the area criteria specified in Step 1. Generation of single-cell images in this step allows for the measurement of morphology measures at an unprecedented cellular resolution from hundreds to thousands of different microglia cells, each with unique identifiers.

The single-cell images are used as input for Step 4, which uses Skeletonize (2D/3D) (Lee et al., 1994) and AnalyzeSkeleton (Arganda-Carreras et al., 2010) to generate measures of different morphology features including maximum branch length, average branch length, and numbers of end point voxels, junction voxels, triple points, branches, junctions, slab voxels, and quadruple points for every cell. MicrogliaMorphology saves these outputs as individual .csv files, which contain all skeleton measures for every individual cell in the dataset, marked by unique identifiers. Step 5 involves FracLac (Karperien, A., 1999), a plugin separate from MicrogliaMorphology which uses fractal analysis to measure additional morphology features including the width of bounding rectangle, maximum radius from hull’s center of mass, maximum span across hull, diameter of bounding circle, maximum radius from circle’s center of mass, perimeter, mean radius, mean radius from circle’s center of mass, area, foreground pixels, height of bounding rectangle, max/min radii from circle’s center of mass, relative variation (CV) in radii from circle’s center of mass, span ratio of hull (major/minor axis), max/min radii from hull’s center of mass, relative variation (CV) in radii from hull’s center of mass, density of foreground pixels in hull area, and circularity. Because FracLac is incompatible with the IJM language, it was not integrated into our MicrogliaMorphology macro and Step 5 must be completed using FracLac-specific user prompts and common parameters outlined in (Young & Morrison, 2018). Steps to batch process the single-cell images generated from MicrogliaMorphology using FracLac are written out in detail on our Github page for MicrogliaMorphology. Importantly, the unique identifiers for each cell are retained in the FracLac output, allowing for integration with the MicrogliaMorphology measures. The final AnalyzeSkeleton output from MicrogliaMorphology (Step 4) and FracLac (Step 5) are merged using MicrogliaMorphologyR to generate a final, master .csv file containing measures for 27 different morphology features for every individual cell.

An additional feature within MicrogliaMorphology is the ColorByCluster feature, which allows the user to color the microglia cells in the original .tiff input images by their k-means cluster identification (see MicrogliaMorphologyR section below). This is a unique feature of MicrogliaMorphology that allows the user to visually validate their morphological clusters and gain insight into their spatial distribution in the brain. (Fig. 3E) All original input immunofluorescent .tiff images containing FASTMAP subregion ROIs, thresholded images generated by MicrogliaMorphology, thresholded single cell images generated by MicrogliaMorphology, and all analysis code for the data presented in this paper will be available on the Open Science Framework (OSF) when the paper is published. Final, tidied up datasets that were used for analysis in this paper are included as part of MicrogliaMorphologyR and can be loaded with the package (data_1xLPS_mouse, data_2xLPS_mouse, data_2xLPS_mouse_fuzzykmeans, and data_ImageTypeComparison).

### MicrogliaMorphologyR

MicrogliaMorphologyR is an R package that wraps several existing packages including tidyverse, Hmisc, pheatmap, factoextra, lmerTest, lme4, Matrix, SciViews, ggpubr, glmmTMB, DHARMa, rstatix, and gridExtra. (Auguie, B., 2017; Bates, D. et al., 2023; Bates et al., 2015; Brooks et al., 2017; Grosjean, P., 2022; Harrell Jr F, 2023; Hartig, F., 2022; Kassambara, A, 2020; Kassambara, A., 2023a, 2023b; Kolde R, 2019; Kuznetsova et al., 2017; Wickham et al., 2019) While our ImageJ macro, MicrogliaMorphology, facilitates the semi-automated measurement of 27 individual morphology features at a single-cell level, our complementary R package, MicrogliaMorphologyR, allows for analysis and visualization of this data to characterize microglia morphological states and gain insight into their relevance in experimental models. Functions within MicrogliaMorphologyR integrate correlation analyses and statistical modeling approaches and are used in conjunction with dimensionality reduction by principal components analysis and k-means clustering to characterize morphological states and quantify population shifts in the experimental model of choice. MicrogliaMorphologyR also includes exploratory data analysis functions to generate heatmap and boxplot visualizations of data in flexible ways including at the single-cell level, animal-level, and experimental condition-level. Furthermore, MicrogliaMorphologyR includes functions for generating quality control metrics on input data such as identifying values that dominate and disproportionately skew feature distributions, data normalization options, and performing linear mixed effects modeling, ANOVA, and other statistical analyses on the input dataset. All source code for MicrogliaMorphologyR and descriptions of functions can be found on Github at: https://github.com/ciernialab/MicrogliaMorphologyR.

## 3 Results

### Morphology analysis toolset: ImageJ macro MicrogliaMorphology

Using ImageJ plugins (Karperien, A., 1999; Young & Morrison, 2018), we have developed an accessible and user-friendly ImageJ tool, MicrogliaMorphology, that automates the characterization of a vast range of morphology features from hundreds to thousands of individual microglial cells. (Fig. 1A) We compiled the ImageJ code into an ImageJ macro format such that users can simply click through options for their morphology analysis and specify where to read and write files. The following steps are automated within the MicrogliaMorphology ImageJ macro such that users can easily perform morphology analysis by following user prompts within ImageJ. Briefly, immunofluorescence images are binarized, cleaned up to remove any background noise, and thresholded according to standard protocol (Young & Morrison, 2018) before individual cell images are created based on area measurements to exclude any artifacts that arise as a product of 2D image representation. The newly generated single-cell images are then used downstream as input for ImageJ plugins AnalyzeSkeleton (Arganda-Carreras et al., 2010) and FracLac (Karperien, A., 1999; Young & Morrison, 2018) (Fig. 1A) to measure 27 unique morphology features from individual cells in a high-throughput and semi-automated manner. (Fig. 1A) Importantly, MicrogliaMorphology saves the ROI coordinate information of each individual cell in the output file so that the user has the option of linking the individual cells back to their spatial locations in the original input images. We have enabled this option through the complementary ColorByCluster functions in MicrogliaMorphology and MicrogliaMorphologyR.

As MicrogliaMorphology is wrapped around the ImageJ plugin FracLac (Karperien, A., 1999), which can only handle 2D input, it is limited to the analysis of 2D images. While 3D reconstructions of microglia offer the benefit of finer-grained detail for morphology analysis, the size of the generated datasets and the time and resources necessary to construct and process such data limits the applicability of 3D measurements to smaller areas, inevitably making the analysis comparatively low-throughput. 2D image options include maximum projection Extended Depth of Focus (EDF) images, which only compile the pixels of highest intensity at any position in a z-stack to construct a new 2D image which preserves some 3D information, vs. individual 2D images sampled that only capture single-plane information. To assess how accurately microglial morphology is represented in 3D image forms as compared to 2D image forms (EDF, single-plane 2D), we analyzed just the skeletal measures (Fig. 2E) from 20 individual microglial cells manually isolated from z-stack images of the hippocampus of female mice. AnalyzeSkeleton, one of the ImageJ plugins that MicrogliaMorphology is wrapped around, is able to measure skeletal features in both 2D and 3D, enabling direct comparisons across different image forms. 5 cells of each morphological type (ameboid, ramified, hypertrophic, rod-like) were manually classified and selected out from the original 3D z-stacks, from which EDF images were generated and 2D single-plane images from the center of the stack were saved to create the test dataset.

**Figure 2:**
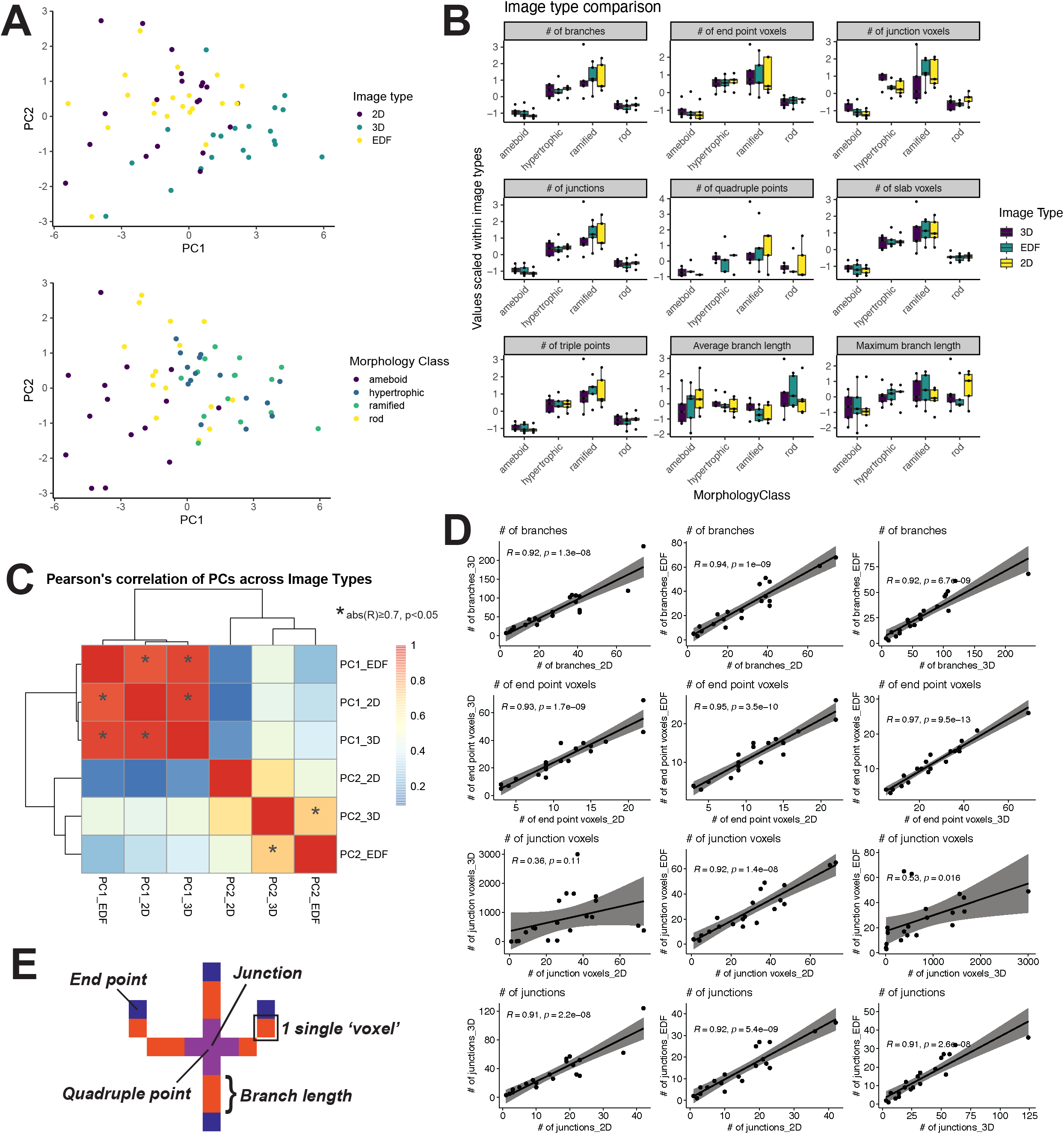
Comparison of 3D vs. Extended Depth of Focus vs. Single-plane 2D image types. (A) Samples represented in Principal Components space and colored by image type or morphology class. Each point is either a 2D, 3D, or EDF representation of one of twenty different cells. (B) Comparison of changes across morphological classes when cells are represented in 2D, 3D, or EDF forms. Values on plots are z-scores (centered and scaled) calculated within image type. (C) Spearman’s correlation of PCs 1-2 after dimensionality reduction across image types. (D) Individual Pearson correlations between image types for specific morphology features measured using AnalyzeSkeleton. (E) Visual description of morphology features measured using AnalyzeSkeleton.

**Figure 3:**
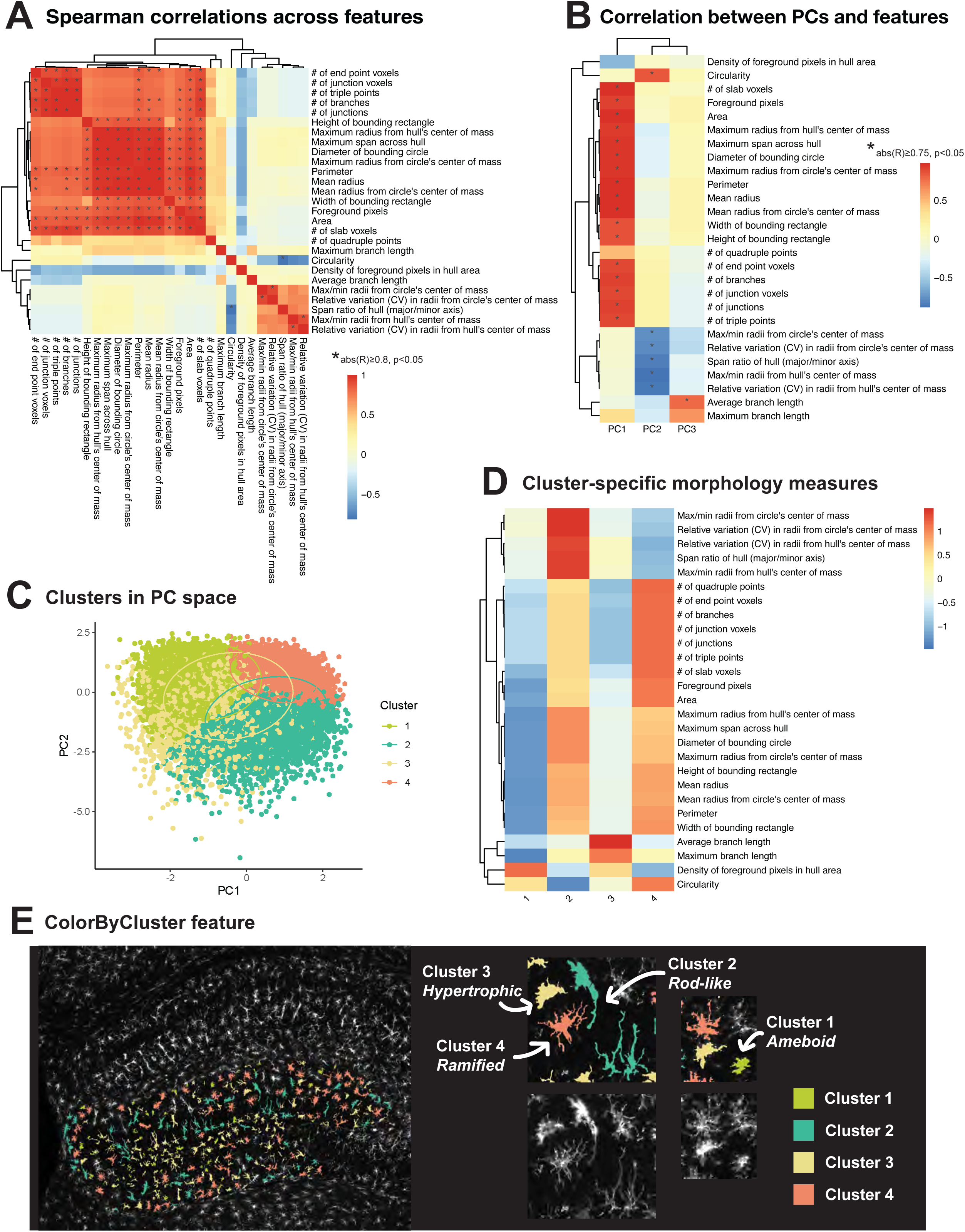
Characterization of morphological clusters in 2xLPS dataset. (A) Spearman’s correlation matrix of 27 features measured by MicrogliaMorphology. (B) Spearman’s correlation of morphology measures to first 3 PCs after dimensionality reduction. (C) Cluster classes displayed in PCs 1-2 space. (D) Average values for all 27 morphology features, scaled across clusters. (E) Individual cells spatially registered back to original images and visually annotated by morphological class using ColorByCluster feature.

As expected, the raw values of all skeletal measures quantified decreased considerably when measured from 3D to either of the 2D image forms (Supp. Fig. 1A). While cells with more ramification and cell branching complexity (hypertrophic, ramified classes) were better captured in 3D (Fig. 2A), relative differences in morphology across the four different forms were well conserved across image types (Fig. 2B). After dimensionality reduction, PC2, which mostly captured variability described by the maximum branch length (Supp. Fig 1C), was highly correlated between 3D and EDF forms but not 3D and 2D forms (Fig. 2C, Supp. Fig. 1B). Cell branching complexity, as described by numbers of junctions, end point voxels, branches, slab voxels, and triple points, was highly correlated across image types (Fig. 2D, Supp. Fig. 1B). Although the number of junctions was also highly correlated across image types, junction voxels, or the numbers of actual pixels which make up the junctions, had low correlation scores when comparing 3D images with either of the 2D image types (Fig. 2D), which is unsurprising, as a 3D image would retain much more of this kind of information. Numbers of quadruple points, which would describe the most complex type of junction, was the least well-captured in 2D representations (Supp. Fig. 1B). As expected, these results together indicate that EDF images better retain 3D skeletal information than 2D forms do, but that relative differences between ameboid, hypertrophic, ramified, and rod-like morphologies are still maintained across image forms.

### Morphology analysis toolset: R package MicrogliaMorphologyR

Once the 27 morphological features are measured from individual microglia using the MicrogliaMorphology ImageJ macro and FracLac, they are concatenated into a final output data file which can be analyzed further to gain insight into microglia morphology changes in any given experimental model. We have provided an R package, MicrogliaMorphologyR, which contains a set of functions that implement one set of approaches for such an analysis in R. Using MicrogliaMorphologyR, the user can conduct exploratory data analysis to generate visualizations of their own data in flexible ways including heatmaps of how morphological features vary across morphological clusters and boxplots of how morphological populations shift at the subject-level across treatment conditions. Functions within MicrogliaMorphologyR are used in conjunction with principal component analysis and k-means clustering to gain further insight into microglia morphology features, classify individual cells by their morphological states, and allow for the quantification of morphological population shifts in experimental contexts. MicrogliaMorphologyR also includes functions for generating quality control metrics on input data such as identifying values that dominate and disproportionately skew feature distributions, data normalization options, visiually exploring different sources of variability in the dataset, and performing ANOVA analysis, linear mixed effects modeling, and other statistical analyses on the input dataset.

### Application to 2xLPS mouse dataset

To demonstrate the utility of MicrogliaMorphology and MicrogliaMorphologyR, we describe an experimental dataset collected from the brains of 8-week old Cx3cr1-GFP mice. Male and female mice were injected peripherally with 2 daily injections of vehicle or 0.5 mg/kg lipopolysaccharide (LPS), a major structural component of gram-negative bacteria that is commonly used to induce and study microglial responses in the brain. (Fig. 1B) To capture a diverse range of microglia morphologies across multiple brain regions, we focused on profiling the frontal cortex, striatum, and hippocampus from 6 individual mice (n=3/treatment, n=2 females and 1 male/treatment). Using MicrogliaMorphology, we were able to quantify 27 morphological features from a total of 43,332 individual microglial cells, which made up our input dataset for analysis.

Within the pool of morphology measures, we defined distinct sets of highly correlated features that describe different aspects of morphology, including branching complexity, area and territory span, branch length, and cell shape. (Fig. 3A) Spearman correlation analysis across all 27 features revealed that those which describe branching complexity (number of end point voxels, junction voxels, triple points, branches, junctions) were highly correlated to each other compared to the other features (R≥0.8; p<0.05). Similar correlations were observed for features that describe area and territory span (width of bounding rectangle, maximum radius from hull’s center of mass, maximum span across hull, diameter of bounding circle, maximum radius from circle’s center of mass, perimeter, mean radius, mean radius from circle’s center of mass, area, number of slab voxels, foreground pixels, height of bounding rectangle), branch length (maximum branch length, average branch length), and cell shape (max/min radii from bounding circle’s and hull’s center of mass, relative variation in radii from bounding circle’s and hull’s center of mass). As expected, cell circularity was highly negatively correlated (R≤-0.8; p<0.05) with span ratio of the bounding hull (major/minor axis), a measure whose higher value indicates greater cell oblongness (Fig. 3A). The relationships observed among the 27 morphology features were consistently captured in another LPS dataset collected under entirely different conditions (reversed light cycle, single 1.0 mg/kg LPS exposure, 24 hour collection time) (Supp. Fig. 3A)., demonstrating that MicrogliaMorphology is able to consistently and robustly capture different aspects of microglia morphology across experimental models.

### Dimensionality reduction and soft clustering

To define morphological states from our 27-feature dataset, we performed dimensionality reduction using principal component analysis followed by fuzzy k-means clustering on the first three principal components (PCs), which together explained 84.6% of the variability in the dataset (Supp. Fig. 2B). Spearman’s correlation of the first 3 PCs to the 27 features showed that each PC was differentially correlated to and described by different sets of morphology features (abs(R)≥0.75; p<0.05) (Fig. 3B). PC1 was highly positively correlated to features describing branching complexity and territory span, meaning that individual cells with greater branching complexity or area had higher PC1 scores (Fig. 3B). PC1 was also highly positively correlated to density of foreground pixels in hull area, which describes a cell’s occupancy within its territory and can be a proxy for soma and/or branch thickness. Taking these correlations together, PC1 captured the variability in the dataset driven by branching complexity, territory span, and territory occupancy. In a similar manner, PC2 captured variability driven by cell circularity and cell shape and PC3 captured variability driven by average branch length (Fig. 3B). In line with our feature analysis (Fig. 3A, Supp. Fig. 3A), we also observed that the PCs were similarly described by the same distinct sets of features in the 1xLPS dataset (Supp. Fig. 3B).

The first three PCs were used as input downstream for fuzzy k-means clustering (Cebeci, Z., 2019), a soft clustering method that is similar in concept and algorithm to k-means clustering, which partitions data points within a given dataset into defined numbers of clusters based on their proximity to the nearest cluster’s centroid. In fuzzy k-means, data points are not exclusively assigned to just one cluster, but rather given scores of membership to all clusters, allowing for ‘fuzziness’ or overlap between two or more clusters. This allows for additional characterization of high-scoring cells within each cluster, cells with more ambiguous identities, and other cases that the user might be interested in, which might be informative to their specific dataset. Fuzzy k-means also assigns a final ‘hard’ cluster assignment based on the class with the highest membership score, which can be used as input for downstream analysis. These final cluster assignments were then used for the analysis of the 2xLPS mouse dataset in this paper, unless otherwise specified. Using exploratory data analysis methods including the within sum of squares and silhouette methods (Supp. Fig. 2C), we found that a clustering parameter of 4 yields the highest degree of within-cluster similarity and was thus the most optimal parameter to use for our example 2xLPS dataset.

### Cluster characterization and analysis

Once cluster membership was defined using k-means clustering, we further explored what features describe the different clusters (Fig. 3D) and how cells belonging to each cluster visually look using the ColorByCluster feature in MicrogliaMorphology (Fig. 3E). We recommend that users always perform these steps in addition to the initial clustering optimization steps (Supp. Fig. 2C) to verify that the clusters defined within their datasets are morphologically distinct and in line with expected differences in microglia morphology. We computed the average values for all 27 morphology measures, scaled across clusters, to characterize how each morphological cluster was differentially defined by the various morphology measures relative to the other clusters (Fig. 3D). Cluster 1 had the lowest branching complexity and territory span, resembling the classic ameboid shape in the original images upon visual confirmation using the ColorByCluster feature in MicrogliaMorphology. Cluster 2 had the greatest oblongness and branching inhomogeneity, and resembled rod-like shapes; Cluster 3 had the highest branch lengths and density of foreground pixels in the hull with average territory span values relative to the other clusters and appeared hypertrophic; and Cluster 4 had the greatest branching complexity, territory span, and circularity, and appeared ramified. (Fig. 3D, Fig. 3E) Clusters 1 (ameboid), 2 (rod-like), and 4 (ramified) cells had relatively lower overlap in PC space with each other compared to Cluster 3 (hypertrophic) cells, which highly overlapped with Cluster 1 and Cluster 2 cells (Fig. 3C). This was expected, as hypertrophic cells represent a state between ameboid and rod-like forms on the morphological spectrum. (Fig. 3C, Fig. 3E). Clusters were similarly described by the different morphology measures in the independent 1xLPS dataset (Supp. Fig. 3C-D), further pointing to MicrogliaMorphology and MicrogliaMorphologyR as a robust means to characterize and analyze microglia morphologies.

### Analysis of different microglia markers

To test for LPS-induced morphological population shifts at the subject-level, we first calculated the percentage of cells in each morphology cluster for every brain region and antibody separately for every mouse using the ‘clusterpercentage’ function within MicrogliaMorphologyR. To assess how cluster membership changes with LPS treatment across brain regions, we fit a generalized linear mixed model using a beta distribution to model the percentage of cluster membership as a factor of Cluster identity, Treatment, and BrainRegion interactions with Antibody as a fixed effect and MouseID as a repeated measure (“percentage ∼ Cluster*Treatment*BrainRegion + Antibody + (1|MouseID)”) using the ‘stats_cluster.animal’ function from MicrogliaMorphologyR, which is wrapped around the glmmTMB R package (Brooks et al., 2017). (Supp. Info. 2) The beta distribution is suitable for values like percentages or probabilities that are constrained to a range of 0-1. 2-way Analysis of Deviance (Type II Wald chisquare tests) on the model revealed a main effect for Cluster, Treatment, and BrainRegion interactions, X^2^(6, n=6)=20.479, Pr(>Chisq)=0.002. There was no significant effect of Antibody (X^2^(2, n=6)=0.085, Pr(>Chisq)=0.959), and we analyzed the Iba1, Cx3cr1, and P2ry12-stained cells as 3 separate datasets. We first filtered for each individual antibody before fitting updated models using the ‘stats_cluster.animal’ function (“percentage ∼ Cluster*Treatment*BrainRegion + (1|MouseID)”) for each antibody separately. T-tests between Treatments (PBS vs. 2xLPS) were corrected for multiple comparisons across Clusters and BrainRegions using the Bonferroni method (significance at p<0.05, Bonferroni). Using our toolset, we were able to characterize morphological population shifts across brain regions using different microglial markers in our experimental mouse model. Across the frontal cortex, hippocampus, and striatum, LPS-induced changes in morphological cluster membership were more similar between Cx3cr1 and Iba1-stained datasets, compared to changes in the P2ry12-stained dataset. (Fig. 4A) As one example, in the frontal cortex for both Cx3cr1 and Iba1 datasets, the percentage of ameboid and ramified cells significantly decreased while the percentage of hypertrophic cells increased and there was no significant change in the proportion of rod-like cells, indicating a shift towards a hypertrophic state. (Fig. 4A, Supp. Info. 2) In the frontal cortex for the P2ry12 dataset, ameboid cells decreased and hypertrophic cells increased, while there was no significant change in the proportions of ramified and rod-like cells between treatments. (Fig. 4A) Antibody-specific differences were also apparent upon examination of the immunofluorescent images for each of the antibodies, where in the baseline PBS condition, P2ry12 distribution was less concentrated in the cell bodies and more spread throughout the cell branches. (Fig. 4B)

**Figure 4:**
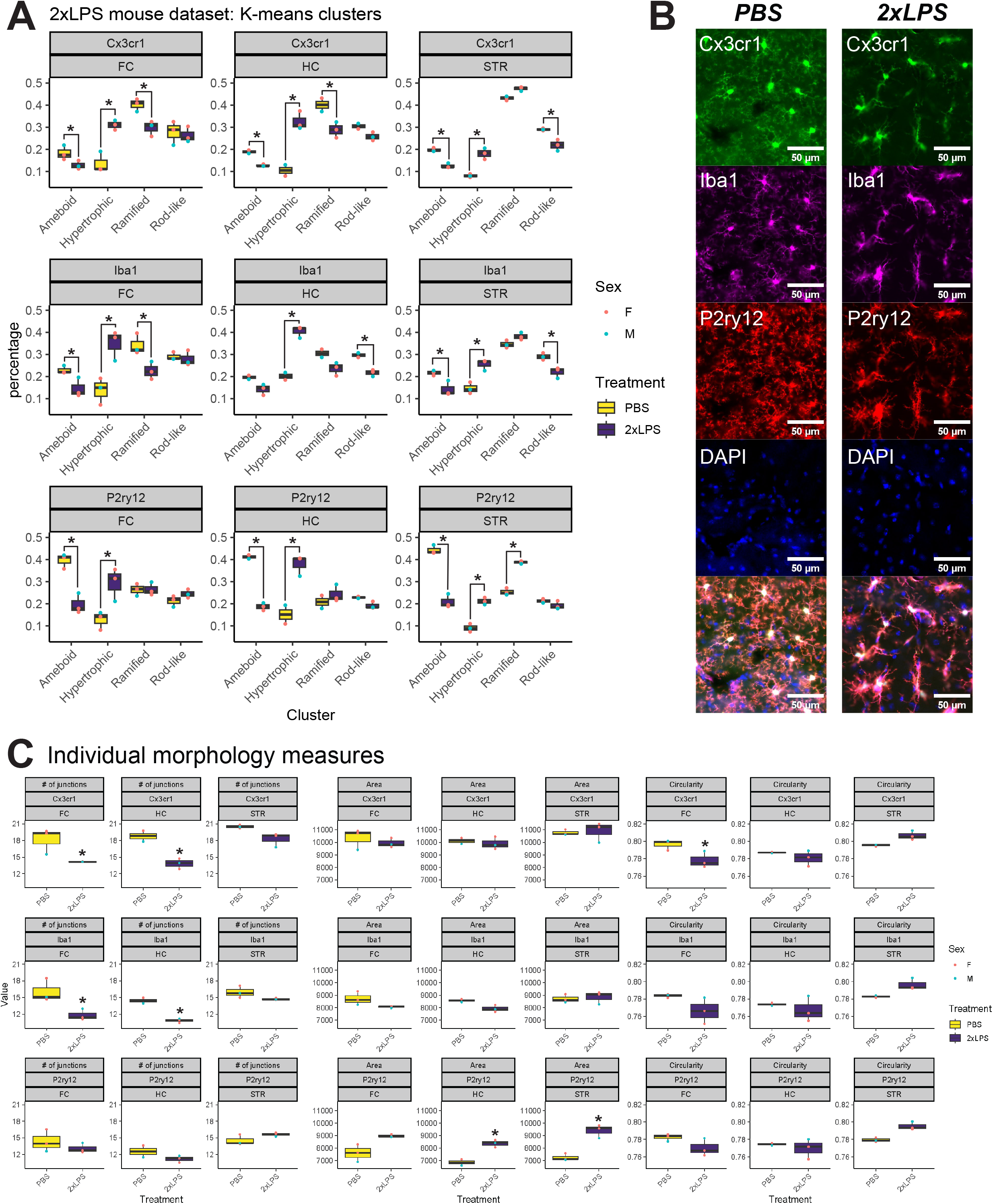
Analysis of morphological clusters and individual morphology measures across brain regions and antibody markers in 2xLPS dataset. (A) LPS-induced shifts in morphological populations across brain regions and antibodies. (*p<0.05, Bonferroni) (B) Immunofluorescent images of the same microglial cells stained with Cx3cr1, Iba1, and P2ry12 in PBS and 2xLPS conditions. Scale bars are 50um (C) LPS-induced changes in individual morphology measures (number of junctions, area, circularity) across brain regions and antibodies. (*p<0.05, Bonferroni)

We also assessed how the 27 individual morphology measures change with LPS treatment by fitting a linear model to the measure values as a factor of Treatment and BrainRegion interactions with Antibody as a fixed effect (“Value ∼ Treatment*BrainRegion + Antibody”). We fit this model for each morphology measure individually using the ‘stats_morphologymeasures.animal’ function from MicrogliaMorphologyR, which is wrapped around the ‘lm’ function in R. We analyzed the Iba1, Cx3cr1, and P2ry12-stained cells as 3 separate datasets (Supp. Info. 2). T-tests between Treatments (PBS vs. 2xLPS) were corrected for multiple comparisons across BrainRegions using the Bonferroni method (significance at p<0.05, Bonferroni). Using our toolset, we were able to characterize changes in specific morphology measures across brain regions and microglial markers. Similar to the changes seen when analyzing LPS-induced shifts in morphological clusters, LPS-induced changes in individual morphology measures were more similar between Cx3cr1-stained and Iba1-stained cells than with P2ry12-stained cells. (Fig. 4B-C) From the 27 measures, we highlight 3 here – the number of junctions, area, and circularity of the cells. (Fig. 4C, Supp. Info. 2) Changes in cell circularity were maintained across all three microglial markers. P2ry12-stained cells showed LPS-induced increases in cell area in the hippocampus and striatum that were not evident in the Iba1 and Cx3cr1-stained cells. LPS-induced decreases in the number of junctions in the frontal cortex and hippocampus were consistent between only the Cx3cr1 and Iba1-stained datasets, while P2ry12-stained cells showed no differences across all brain regions. Taken together, our findings from both the cluster and morphology measure analyses show that Cx3cr1 and Iba1 are more consistent with each other than with P2ry12. Thus, it is important to keep these differences in mind when choosing microglia markers for morphology experiments, and to keep choices consistent within an experiment to avoid antibody-related artifacts and false positives.

## 4 Discussion

### MicrogliaMorphology and MicrogliaMorphologyR, a high-throughput pipeline to characterize microglia morphological states at a single-cell resolution

Microglia exhibit a dynamic range of morphologies including ramified, ameboid, rod-like, and hypertrophic forms that are highly context-specific and often rapidly changing in response to local environmental cues. (Paolicelli et al., 2022; Reddaway et al., 2023; Savage et al., 2019) There has been a concerted effort as a field to move away from dualistic characterization of all microglia as ‘resting’ or ‘activated’, which is often described in terms of morphological differences, and towards a clearer understanding and appreciation for the heterogenous ‘states’ of microglia that co-exist in the brain in any given context. (Dubbelaar et al., 2018; Paolicelli et al., 2022) In line with these efforts, there have been many recently published tools that classify and analyze microglia morphological subpopulations in an automated and high-throughput manner. (Clarke et al., 2021; Colombo et al., 2022; Hrj et al., 2023; Leyh et al., 2021; Reddaway et al., 2023; Salamanca et al., 2019; York et al., 2018) Using our toolset, MicrogliaMorphology and MicrogliaMorphologyR, we take a data-informed approach to characterize different populations of microglia morphologies and to statistically model how membership across all morphological states dynamically changes in experimental contexts and across brain regions in an automated and high-throughput manner, which offers a great advantage over more labor-intensive morphological approaches which employ manual categorizations of cells or assessment of individual measures of morphology rather than morphological states. (Torres-Platas et al., 2014; Young & Morrison, 2018) Furthermore, the ColorByCluster feature within MicrogliaMorphology and functions within MicrogliaMorphologyR facilitate comparisons of morphological measures across clusters and together provide a thorough validation for verifying cluster identities both visually and analytically compared to existing tools. We demonstrate that MicrogliaMorphology and MicrogliaMorphologyR are able to reproducibly detect both subtle and pronounced changes in microglia morphology and together provide a robust method to characterize morphological states across a wide range of experimental and disease models. While our dataset was too underpowered to quantify sex differences in morphology, our tool could be used to explore known sex differences in microglia in various contexts such as early brain development. (Sullivan & Ciernia, 2022)

Importantly, we made both MicrogliaMorphology and MicrogliaMorphologyR free and open source resources. Our toolset only relies on software that is open source and freely available to download and all relevant materials including input images, data, and supporting code used in this study will be available on the OSF website when the paper is published. We will also include all of the single-cell images that were generated for this study at the OSF link, which provides a unique, benchmarking dataset for researchers interested in applying other approaches such as machine learning methods to classify microglia morphology. The ImageJ and R code underlying both MicrogliaMorphology and MicrogliaMorphologyR are all available through Github repositories at https://github.com/ciernialab/MicrogliaMorphology and https://github.com/ciernialab/MicrogliaMorphologyR in the hopes that the larger research community can openly share troubleshooting tips, benefit from discussion, and continue to expand upon our work and develop our toolsets for broader use.

### Choice of markers affects morphology analysis

Cx3cr1, Iba1, and P2ry12 are all antibody markers that are commonly used to visualize and study various aspects of microglia including morphology. Of these 3 markers, P2ry12 is the most microglia-specific, as both Cx3cr1 and Iba1 also label other macrophages. (Paolicelli et al., 2022) While all three of these markers can label microglia reliably in homeostatic conditions and are considered ‘homeostatic’ markers, their expression can change in disease-associated and inflammatory states. For instance, in a study (Kenkhuis et al., 2022) of co-expression patterns of microglia markers Iba1, P2ry12, and Tmem119, another microglia-specific antibody, in the brains of Alzheimer’s Disease patients, P2ry12 expression was lost in microglia surrounding amyloid-beta plaques, while Iba1 expression was increased in subsets of microglia and Tmem119 expression was generally lost across microglia. Peripheral macrophages have also been shown to infiltrate the blood brain barrier and enter the brain in disorders such as Parkinson’s Disease and Multiple Sclerosis (Prinz & Priller, 2017), which could complicate the analysis of morphology in datasets stained with non-specific markers such as Cx3cr1 and Iba1. Thus, careful consideration of morphological markers should be taken depending on the experimental context in which microglia are being studied. (Paolicelli et al., 2022)

In our analyses of LPS-induced shifts in morphological populations and changes in individual morphology measures across three commonly used microglial markers – Cx3cr1, Iba1, and P2ry12 – we found that P2ry12 showed unique differences in the percentage of morphological populations present across brain regions, the directionality of shifts across morphological populations, and the specific morphological features such as the number of junctions and cell area that change with LPS administration. (Fig. 4A, Fig. 4C, Supp. Info. 2) P2ry12 immunoflourescent signal was also more uniformly distributed throughout the entirety of the cell and less localized to the cell soma when compared to Cx3cr1 and Iba1-stained cells (Fig. 4B) (Paolicelli et al., 2022), making P2ry12-stained cells potentially more likely to be recognized as overlapping cells and consequently filtered out based on area during the thresholding and dataset optimization steps in MicrogliaMorphology. Thus, P2ry12-stained datasets may be better suited for analysis in 3D image types. If using MicrogliaMorphology for 2D P2ry12-stained images, the thresholded images and the single-cells extracted from these images should be carefully examined and compared to the original immunofluorescent images to ensure that the cells analyzed are accurately represented before proceeding with analysis and biological interpretation of results.

### Requirements of MicrogliaMorphology and MicrogliaMorphologyR

Our toolset requires the use of 2D image forms (Extended Depth of Focus or single-plane 2D images) and depends on some user input to determine dataset-specific parameters for MicrogliaMorphology. Introductory skills coding in the R language are also necessary to be able to use MicrogliaMorphologyR and larger computational resources may be required for the analysis of larger datasets. However, many of these requirements exist for alternative approaches for morphology analysis that have been presented as well.

### Future directions

To classify microglia by their morphological characteristics in any approach, hard cut-offs are used to define where one class starts and the next begins. This type of binning of morphologies is limited in that microglia are inherently dynamic and more realistically exist along a continuum of morphological forms. To represent the dynamic nature of microglia morphology, we also demonstrate that MicrogliaMorphology and MicrogliaMorphologyR can be optionally integrated with a soft clustering approach using fuzzy k-means clustering, which is similar in concept and algorithm to k-means clustering. Soft clustering approaches such as fuzzy k-means clustering not only yield final cluster (or class) identities for every cell, but also membership scores of belonging to any given cluster. This allows for additional characterization of high-scoring cells within each cluster (i.e., quintessential ‘rod-like’, ‘ameboid’, ‘hypertrophic’, or ‘ramified’ cells), cells with more ambiguous identities (e.g., a cell that is 5% rod-like, 5% ameboid, 45% hypertrophic, and 45% ramified), and other cases that the user might be interested in which might be informative for their specific dataset. Fuzzy k-means also assigns a final hard cluster assignment for every cell based on the class with the highest membership score, so the user can also use these final assignments as input for downstream analysis. While we used the hard cluster assignments for the analysis in this paper, we provide an example of using the soft clustering assignments from fuzzy k-means to analyze just the high-scoring cells for each morphological class at the end of the Github page for MicrogliaMorphologyR. While we used k-means clustering approaches in this study, our toolset is highly flexible and can also be integrated with other clustering approaches such as hierarchical clustering or gaussian mixture models.

Microglia have long been known as a highly heterogenous cell type as defined by their morphology, electrophysiological properties, transcriptomic profiles, and surface expression of immune markers. (Hammond et al., 2019; Li et al., 2019; Masuda et al., 2020; Paolicelli et al., 2022) Context-specific regulation of morphology further emphasizes the need to probe microglial phenotypes from multiple angles in conjunction with morphology to gain more clarity on the relationship between microglial form and function. (Dubbelaar et al., 2018; Paolicelli et al., 2022) While the majority of studies of microglia morphology have yielded observational insights into the range of forms present in various contexts, only a few (Adrian et al., 2023; Madry et al., 2018, p. 1; Parakalan et al., 2012) have actually explored how different morphological states directly contribute to microglial function in the brain. The rise of single-cell sequencing technologies has provided vast new insight into the molecular mechanisms that shape heterogenous microglial responses and has granted us a better understanding of microglial ‘states’ in homeostatic, developmental, and disease-relevant contexts (Hammond et al., 2019; Li et al., 2019; Masuda et al., 2020). However, transcriptomic characterization alone does not capture the diversity of changes that microglia exhibit and we still lack a direct understanding of whether morphologically different microglia populations are transcriptomically distinct. (Parakalan et al., 2012) is one such study that directly explored these relationships by identifying over 2000 differentially expressed genes with unique sets of biological functions between ameboid and ramified microglia laser-dissected and pooled from rat brains. However, while it’s agreed upon that microglia exhibit a wide range of morphological forms across various biological contexts, it is still unclear whether we can transcriptomically define the heterogeneous morphological states that exist outside of the ‘resting’ vs. ‘activated’ morphological dichotomy and in what ways these transcriptomic signatures relate to microglial function.

The advent of spatially-resolved transcriptomics and development of methods for integrating multiple data modalities has opened new avenues to explore these relationships more directly. The ability to map morphologically-classified microglia back to their spatial locations in their original input images using the complementary ColorByCluster functions in MicrogliaMorphology and MicrogliaMorphologyR allows for not only the visual verification and exploration of morphological cluster identity across tissue sections, but also facilitates the direct integration of spatial transcriptomics data to morphological data at a cellular resolution. Our toolset serves as a resource that can complement new tools and approaches such as spatial transcriptomics to answer questions about the relationship between microglia morphology and microglia function more directly.

## Acknowledgements

We thank the Ciernia Lab and Pavlidis Lab members for their thoughtful feedback and suggestions during lab meetings throughout the progression of this project. We would also like to thank Wai Hang (Tom) Cheng, whose help was instrumental with learning how to image on the Axioscan slidescanner and getting started with microglia morphology analysis; Nicholas Michelson, whose help was invaluable when troubleshooting code in ImageJ for various features of MicrogliaMorphology; and Dylan Terstege, who generously provided the materials for FASTMAP alignment to the Allen Brain Atlas before they were published. We would also like to thank Dr. Brian MacVicar for sharing his lab’s Cx3cr1-GFP mice with us, which we used for the 2xLPS *in vivo* experiments. We are grateful for the computational resources provided by the Neuroimaging & Neurocomputation Centre and the Dynamic Brain Circuits in Health & Disease Research Cluster at the University of British Columbia. This work was supported by the Canadian Open Neuroscience Platform Student Scholar Award (10901 to JK); University of British Columbia Four Year Doctoral Fellowship (6569 to JK); Canadian Institutes for Health Research (CRC-RS 950-232402 to AC); Natural Sciences and Engineering Research Council of Canada (RGPIN-2019-04450, DGECR-2019-00069 to AC); Scottish Rite Charitable Foundation (21103 to AC) and the Brain Canada Foundation (AWD-023132 to AC). The funders had no role in study design, data collection and analysis, decision to publish, or preparation of the manuscript.

---

**Supplementary Figure 1:**
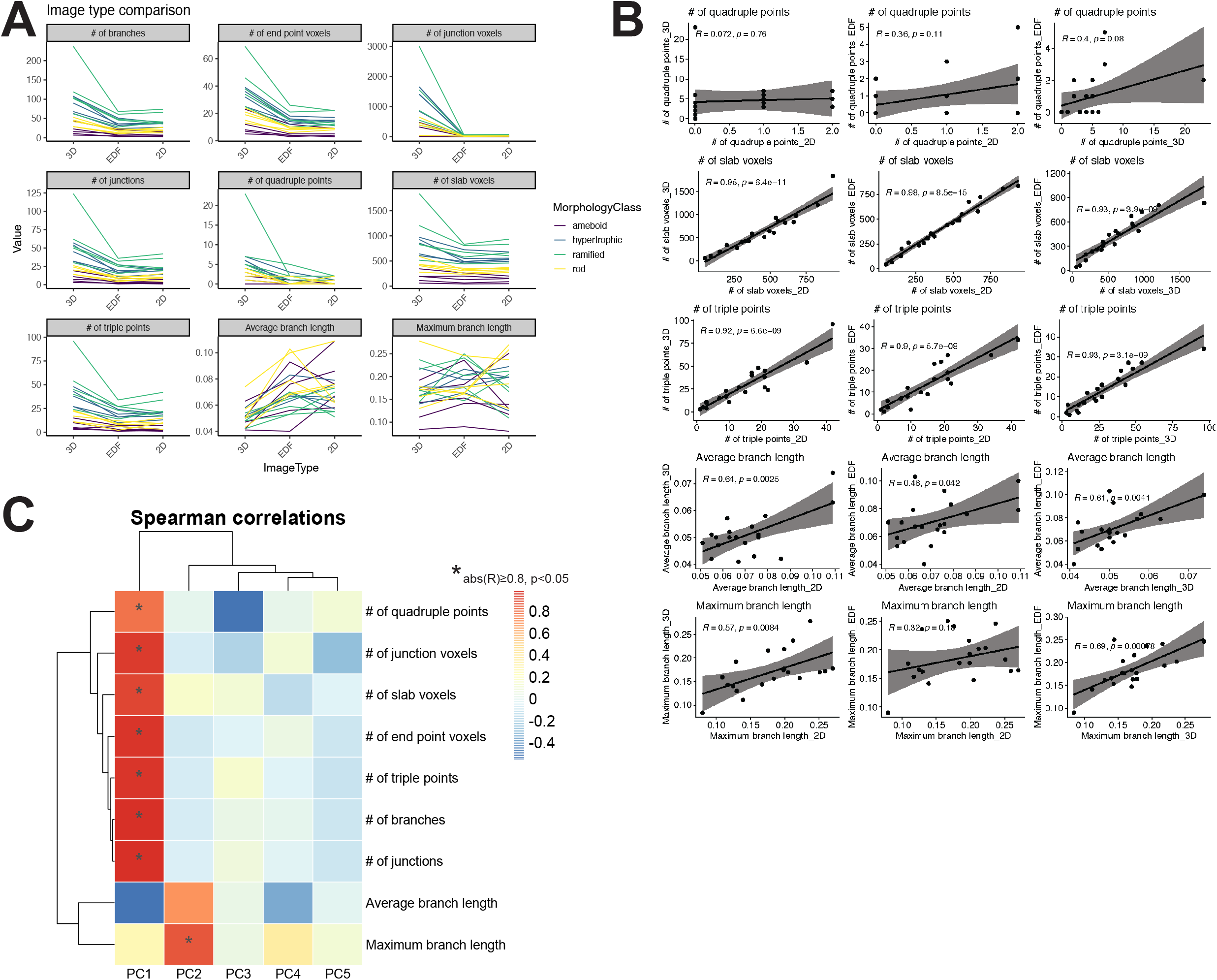
Extended comparison of 3D vs. 2D image types. (A) Changes in the raw values of AnalyzeSkeleton measures across 3D, EDF, and 2D image types. Each line is an individual cell represented in 3D, EDF, and 2D. (B) Individual Pearson’s correlations between image types for specific morphology features measured using AnalyzeSkeleton. (C) Spearman’s correlation of skeletal morphology measures to first 5 PCs after dimensionality reduction.

**Supplementary Figure 2:**
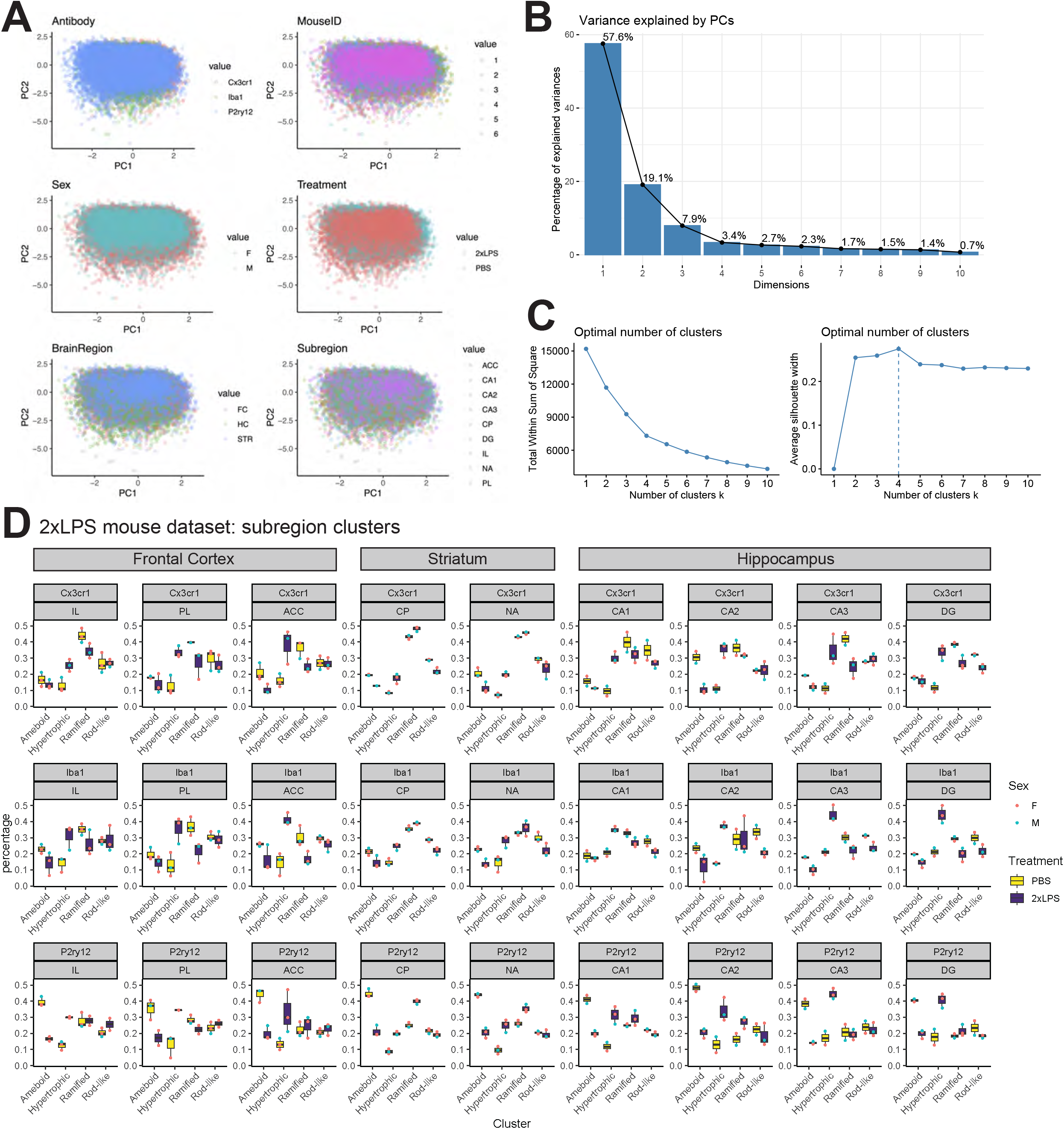
Extended analysis of 2xLPS dataset. (A) Cells from dataset visualized in PCs 1-2 space and colored by different experimental variables. (B) Elbow plot depicting percentage of the variance in dataset explained by each Principal Component. (C) LPS-induced shifts in morphological populations across subregions and antibodies.

**Supplementary Figure 3:**
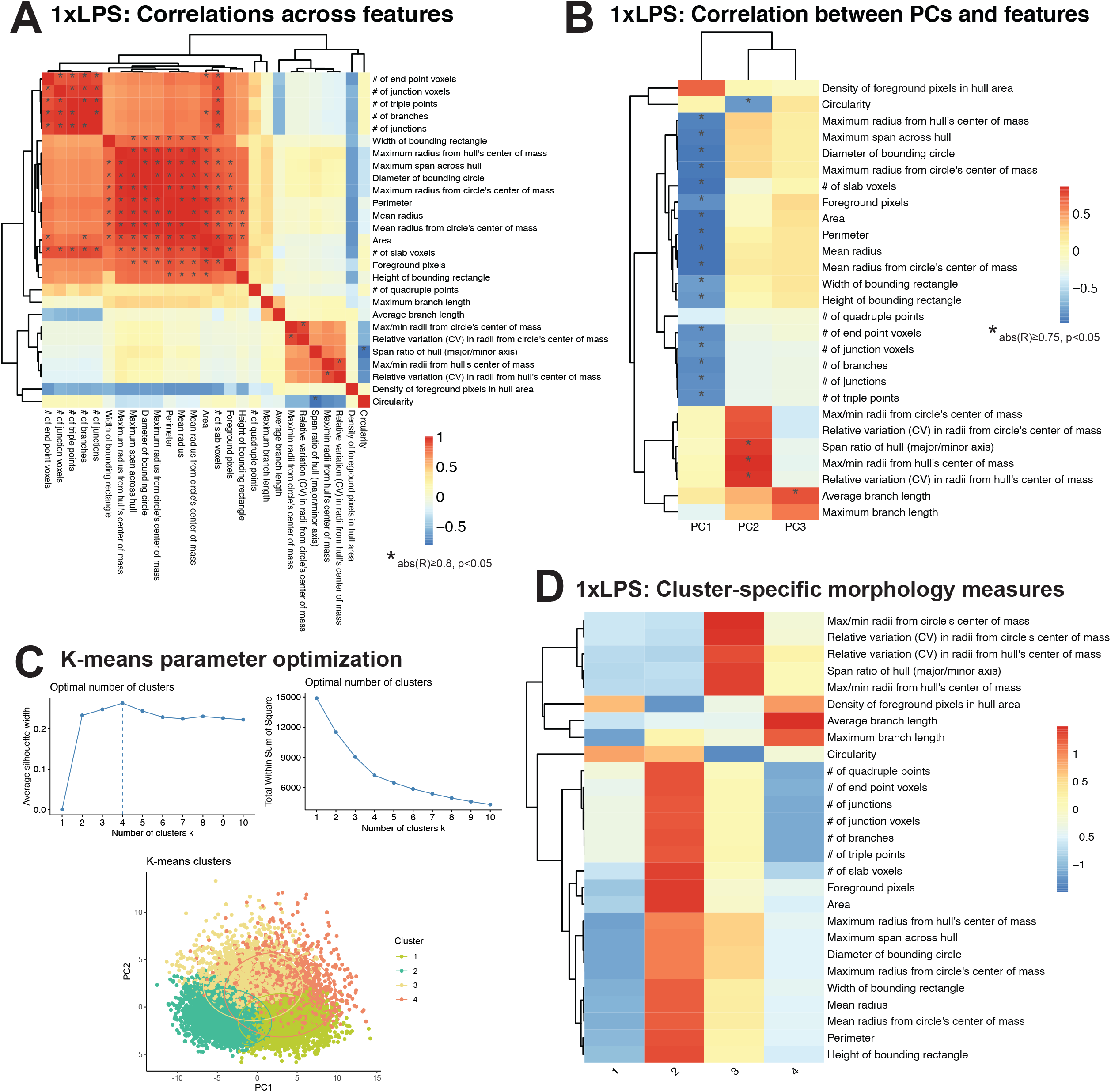
Analysis of 1xLPS morphology measures and clusters. (A) Spearman’s correlation matrix of 27 features measured by MicrogliaMorphology. (B) Spearman’s correlation of morphology measures to first 3 PCs after dimensionality reduction. (C) Optimal k-means clustering parameters determined using within sum of squares and gap statistic techniques. Cluster classes displayed in PC space. (D) Average values for all 27 morphology features, scaled across clusters.

***Supplementary Information 1: Image Type Comparison analysis.*** Plots and underlying code used to generate Figure 2 and Supplementary Figure 1.

***Supplementary Information 2: 2xLPS dataset analysis.*** Plots, statistical analysis and results, and underlying code used to generate Figure 3, Figure 4, and Supplementary Figure 2.

***Supplementary Information 3: 1xLPS dataset analysis.*** Plots and underlying code used to generate Supplementary Figure 3.

## References

Abiega, O., Beccari, S., Diaz-Aparicio, I., Nadjar, A., Layé, S., Leyrolle, Q., Gómez-Nicola, D., Domercq, M., Pérez-Samartín, A., Sánchez-Zafra, V., Paris, I., Valero, J., Savage, J. C., Hui, C.-W., Tremblay, M.-È., Deudero, J. J. P., Brewster, A. L., Anderson, A. E., Zaldumbide, L., … Sierra, A. (2016). Neuronal Hyperactivity Disturbs ATP Microgradients, Impairs Microglial Motility, and Reduces Phagocytic Receptor Expression Triggering Apoptosis/Microglial Phagocytosis Uncoupling. PLOS Biology, 14(5), e1002466. 10.1371/journal.pbio.1002466

Adrian, M., Weber, M., Tsai, M.-C., Glock, C., Kahn, O. I., Phu, L., Cheung, T. K., Meilandt, W. J., Rose, C. M., & Hoogenraad, C. C. (2023). Polarized microtubule remodeling transforms the morphology of reactive microglia and drives cytokine release. Nature Communications, 14(1), Article 1. 10.1038/s41467-023-41891-6

Arganda-Carreras, I., Fernández-González, R., Muñoz-Barrutia, A., & Ortiz-De-Solorzano, C. (2010). 3D reconstruction of histological sections: Application to mammary gland tissue. Microscopy Research and Technique, 73(11), 1019–1029. 10.1002/jemt.20829

Auguie, B. (2017). gridExtra: Miscellaneous Functions for “Grid” Graphics (R package version 2.3) [Computer software]. https://CRAN.R-project.org/package=gridExtra

Bates, D., Mächler, M., Bolker, B., & Walker, S. (2015). Fitting Linear Mixed-Effects Models Using lme4. Journal of Statistical Software, 67, 1–48. 10.18637/jss.v067.i01

Bates, D., Maechler, M., & Jagan, M. (2023). Matrix: Sparse and Dense Matrix Classes and Methods (R package version 1.5-4.1) [Computer software]. https://CRAN.R-project.org/package=Matrix

Bernier, L.-P., York, E. M., Kamyabi, A., Choi, H. B., Weilinger, N. L., & MacVicar, B. A. (2020). Microglial metabolic flexibility supports immune surveillance of the brain parenchyma. Nature Communications, 11(1), Article 1. 10.1038/s41467-020-15267-z

Beynon, S. B., & Walker, F. R. (2012). Microglial activation in the injured and healthy brain: What are we really talking about? Practical and theoretical issues associated with the measurement of changes in microglial morphology. Neuroscience, 225, 162–171. 10.1016/j.neuroscience.2012.07.029

Bialas, A. R., & Stevens, B. (2013). TGF-β signaling regulates neuronal C1q expression and developmental synaptic refinement. Nature Neuroscience, 16(12), 1773–1782. 10.1038/nn.3560

Brooks, M. E., Kristensen, K., Benthem, K. J. van, Magnusson, A., Berg, C. W., Nielsen, A., Skaug, H. J., Mächler, M., & Bolker, B. M. (2017). glmmTMB Balances Speed and Flexibility Among Packages for Zero-inflated Generalized Linear Mixed Modeling. The R Journal, 9(2), 378– 400.

Cebeci, Z. (2019). Comparison of Internal Validity Indices for Fuzzy Clustering | Journal of Agricultural Informatics. Journal of Agricultural Informatics, 10(2), 1–14. 10.17700/jai.2019.10.2.537

Ciernia, A. V., Careaga, M., Ashwood, P., & LaSalle, J. (2018). Microglia from offspring of dams with allergic asthma exhibit epigenomic alterations in genes dysregulated in autism. Glia, 66(3), 505–521. 10.1002/glia.23261

Clarke, D., Crombag, H. S., & Hall, C. N. (2021). An open-source pipeline for analysing changes in microglial morphology. Open Biology, 11(8), 210045. 10.1098/rsob.210045

Colombo, G., Cubero, R. J. A., Kanari, L., Venturino, A., Schulz, R., Scolamiero, M., Agerberg, J., Mathys, H., Tsai, L.-H., Chachólski, W., Hess, K., & Siegert, S. (2022). A tool for mapping microglial morphology, morphOMICs, reveals brain-region and sex-dependent phenotypes. Nature Neuroscience, 25(10), Article 10. 10.1038/s41593-022-01167-6

Cunningham, C. L., Martínez-Cerdeño, V., & Noctor, S. C. (2013). Microglia Regulate the Number of Neural Precursor Cells in the Developing Cerebral Cortex. The Journal of Neuroscience, 33(10), 4216–4233. 10.1523/JNEUROSCI.3441-12.2013

Davalos, D., Grutzendler, J., Yang, G., Kim, J. V., Zuo, Y., Jung, S., Littman, D. R., Dustin, M. L., & Gan, W.-B. (2005). ATP mediates rapid microglial response to local brain injury in vivo. Nature Neuroscience, 8(6), Article 6. 10.1038/nn1472

Dubbelaar, M. L., Kracht, L., Eggen, B. J. L., & Boddeke, E. W. G. M. (2018). The Kaleidoscope of Microglial Phenotypes. Frontiers in Immunology, 9. https://www.frontiersin.org/article/10.3389/fimmu.2018.01753

Geirsdottir, L., David, E., Keren-Shaul, H., Weiner, A., Bohlen, S. C., Neuber, J., Balic, A., Giladi, A., Sheban, F., Dutertre, C.-A., Pfeifle, C., Peri, F., Raffo-Romero, A., Vizioli, J., Matiasek, K., Scheiwe, C., Meckel, S., Mätz-Rensing, K., van der Meer, F., … Prinz, M. (2019). Cross-Species Single-Cell Analysis Reveals Divergence of the Primate Microglia Program. Cell, 179(7), 1609–1622.e16.10.1016/j.cell.2019.11.010

Grosjean, P. (2022). SciViews-R [Computer software]. http://www.sciviews.org/SciViews-R

Hammond, T. R., Dufort, C., Dissing-Olesen, L., Giera, S., Young, A., Wysoker, A., Walker, A. J., Gergits, F., Segel, M., Nemesh, J., Marsh, S. E., Saunders, A., Macosko, E., Ginhoux, F., Chen, J., Franklin, R. J. M., Piao, X., McCarroll, S. A., & Stevens, B. (2019). Single-Cell RNA Sequencing of Microglia throughout the Mouse Lifespan and in the Injured Brain Reveals Complex Cell-State Changes. Immunity, 50(1), 253–271.e6. 10.1016/j.immuni.2018.11.004

Hammond, T. R., Robinton, D., & Stevens, B. (2018). Microglia and the Brain: Complementary Partners in Development and Disease. Annual Review of Cell and Developmental Biology, 34, 523–544. 10.1146/annurev-cellbio-100616-060509

Hansen, D. V., Hanson, J. E., & Sheng, M. (2018). Microglia in Alzheimer’s disease. The Journal of Cell Biology, 217(2), 459–472. 10.1083/jcb.201709069

Harrell Jr F. (2023). Hmisc: Harrell Miscellaneous (R package version 5.1-1) [Computer software]. https://CRAN.R-project.org/package=Hmisc

Hartig, F. (2022). DHARMa: Residual Diagnostics for Hierarchical (Multi-Level / Mixed) Regression Models (R package version 0.4.6) [Computer software]. https://CRAN.R-project.org/package=DHARMa

Hellwig, S., Brioschi, S., Dieni, S., Frings, L., Masuch, A., Blank, T., & Biber, K. (2016). Altered microglia morphology and higher resilience to stress-induced depression-like behavior in CX3CR1-deficient mice. *Brain*, Behavior, and Immunity, 55, 126–137. 10.1016/j.bbi.2015.11.008

Hrj, van W., Tw, N., Ml, B., Ewgm, B., & Bjl, E. (2023). Microglia morphotyping in the adult mouse CNS using hierarchical clustering on principal components reveals regional heterogeneity but no sexual dimorphism. Glia, 71(10). 10.1002/glia.24427

Jung, H., Lee, D., You, H., Lee, M., Kim, H., Cheong, E., & Um, J. W. (2023). LPS induces microglial activation and GABAergic synaptic deficits in the hippocampus accompanied by prolonged cognitive impairment. Scientific Reports, 13(1), Article 1. 10.1038/s41598-023-32798-9

Jung, S., Aliberti, J., Graemmel, P., Sunshine, M. J., Kreutzberg, G. W., Sher, A., & Littman, D. R. (2000). Analysis of fractalkine receptor CX(3)CR1 function by targeted deletion and green fluorescent protein reporter gene insertion. Molecular and Cellular Biology, 20(11), 4106–4114. 10.1128/MCB.20.11.4106-4114.2000

Karperien, A. (1999, 2013). FracLac for ImageJ. https://imagej.nih.gov/ij/plugins/fraclac/FLHelp/Introduction.htm

Kassambara, A. (2020). factoextra: Extract and Visualize the Results of Multivariate Data Analyses (R package version 1.0.7) [Computer software]. https://CRAN.R-project.org/package=factoextra

Kassambara, A. (2023a). ggpubr: “ggplot2” Based Publication Ready Plots (R package version 0.6.0) [Computer software]. https://CRAN.R-project.org/package=ggpubr

Kassambara, A. (2023b). rstatix: Pipe-Friendly Framework for Basic Statistical Tests (R package version 0.7.2) [Computer software]. https://CRAN.R-project.org/package=rstatix

Kenkhuis, B., Somarakis, A., Kleindouwel, L. R. T., van Roon-Mom, W. M. C., Höllt, T., & van der Weerd, L. (2022). Co-expression patterns of microglia markers Iba1, TMEM119 and P2RY12 in Alzheimer’s disease. Neurobiology of Disease, 167, 105684. 10.1016/j.nbd.2022.105684

Kolde R. (2019). pheatmap: Pretty Heatmaps (R package version 1.0.12) [Computer software]. https://CRAN.R-project.org/package=pheatmap

Kuznetsova, A., Brockhoff, P. B., & Christensen, R. H. B. (2017). lmerTest Package: Tests in Linear Mixed Effects Models. Journal of Statistical Software, 82, 1–26. 10.18637/jss.v082.i13

Lee, T.-C., Kashyap, R. L., & Chu, C.-N. (1994). Building skeleton models via 3-D medial surface/axis thinning algorithms. CVGIP: Graphical Models and Image Processing, 56(6), 462–478. 10.1006/cgip.1994.1042

Lenz, K. M., & Nelson, L. H. (2018). Microglia and Beyond: Innate Immune Cells As Regulators of Brain Development and Behavioral Function. Frontiers in Immunology, 9, 698. 10.3389/fimmu.2018.00698

Leyh, J., Paeschke, S., Mages, B., Michalski, D., Nowicki, M., Bechmann, I., & Winter, K. (2021). Classification of Microglial Morphological Phenotypes Using Machine Learning. Frontiers in Cellular Neuroscience, 15, 701673. 10.3389/fncel.2021.701673

Li, Q., Cheng, Z., Zhou, L., Darmanis, S., Neff, N. F., Okamoto, J., Gulati, G., Bennett, M. L., Sun, L. O., Clarke, L. E., Marschallinger, J., Yu, G., Quake, S. R., Wyss-Coray, T., & Barres, B. A. (2019). Developmental Heterogeneity of Microglia and Brain Myeloid Cells Revealed by Deep Single-Cell RNA Sequencing. Neuron, 101(2), 207–223.e10. 10.1016/j.neuron.2018.12.006

Madry, C., Kyrargyri, V., Arancibia-Cárcamo, I. L., Jolivet, R., Kohsaka, S., Bryan, R. M., & Attwell, D. (2018). Microglial Ramification, Surveillance, and Interleukin-1β Release Are Regulated by the Two-Pore Domain K+ Channel THIK-1. Neuron, 97(2), 299–312.e6. 10.1016/j.neuron.2017.12.002

Masuda, T., Sankowski, R., Staszewski, O., & Prinz, M. (2020). Microglia Heterogeneity in the Single-Cell Era. Cell Reports, 30(5), 1271–1281. 10.1016/j.celrep.2020.01.010

Meleady, L., Towriss, M., Kim, J., Bacarac, V., Dang, V., Rowland, M. E., & Ciernia, A. V. (2023). Histone deacetylase 3 regulates microglial function through histone deacetylation. Epigenetics, 18(1), 2241008. 10.1080/15592294.2023.2241008

Nimmerjahn, A., Kirchhoff, F., & Helmchen, F. (2005). Resting microglial cells are highly dynamic surveillants of brain parenchyma in vivo. *Science (New York*, N.Y.), 308(5726), 1314–1318. 10.1126/science.1110647

Paolicelli, R. C., Sierra, A., Stevens, B., Tremblay, M.-E., Aguzzi, A., Ajami, B., Amit, I., Audinat, E., Bechmann, I., Bennett, M., Bennett, F., Bessis, A., Biber, K., Bilbo, S., Blurton-Jones, M., Boddeke, E., Brites, D., Brône, B., Brown, G. C., … Wyss-Coray, T. (2022). Microglia states and nomenclature: A field at its crossroads. Neuron, 110(21), 3458–3483. 10.1016/j.neuron.2022.10.020

Parakalan, R., Jiang, B., Nimmi, B., Janani, M., Jayapal, M., Lu, J., Tay, S. S., Ling, E.-A., & Dheen, S. T. (2012). Transcriptome analysis of amoeboid and ramified microglia isolated from the corpus callosum of rat brain. BMC Neuroscience, 13(1), 64. 10.1186/1471-2202-13-64

Pinskiy, V., Jones, J., Tolpygo, A. S., Franciotti, N., Weber, K., & Mitra, P. P. (2015). High-Throughput Method of Whole-Brain Sectioning, Using the Tape-Transfer Technique. PLOS ONE, 10(7), e0102363. 10.1371/journal.pone.0102363

Prinz, M., & Priller, J. (2017). The role of peripheral immune cells in the CNS in steady state and disease. Nature Neuroscience, 20(2), 136–145. 10.1038/nn.4475

Raj, D. D. A., Jaarsma, D., Holtman, I. R., Olah, M., Ferreira, F. M., Schaafsma, W., Brouwer, N., Meijer, M. M., de Waard, M. C., van der Pluijm, I., Brandt, R., Kreft, K. L., Laman, J. D., de Haan, G., Biber, K. P. H., Hoeijmakers, J. H. J., Eggen, B. J. L., & Boddeke, H. W. G. M. (2014). Priming of microglia in a DNA-repair deficient model of accelerated aging. Neurobiology of Aging, 35(9), 2147–2160. 10.1016/j.neurobiolaging.2014.03.025

Reddaway, J., Richardson, P. E., Bevan, R. J., Stoneman, J., & Palombo, M. (2023). Microglial morphometric analysis: So many options, so little consistency. Frontiers in Neuroinformatics, 17. https://www.frontiersin.org/articles/10.3389/fninf.2023.1211188

Salamanca, L., Mechawar, N., Murai, K. K., Balling, R., Bouvier, D. S., & Skupin, A. (2019). MIC-MAC: An automated pipeline for high-throughput characterization and classification of three-dimensional microglia morphologies in mouse and human postmortem brain samples. Glia, 67(8), 1496–1509. 10.1002/glia.23623

Salvador, A. F., de Lima, K. A., & Kipnis, J. (2021). Neuromodulation by the immune system: A focus on cytokines. Nature Reviews Immunology, 21(8), Article 8. 10.1038/s41577-021-00508-z

Savage, J. C., Carrier, M., & Tremblay, M.-È. (2019). Morphology of Microglia Across Contexts of Health and Disease. *Methods in Molecular Biology (Clifton*, N.J.), 2034, 13–26. 10.1007/978-1-4939-9658-2_2

Schafer, D. P., Lehrman, E. K., Kautzman, A. G., Koyama, R., Mardinly, A. R., Yamasaki, R., Ransohoff, R. M., Greenberg, M. E., Barres, B. A., & Stevens, B. (2012). Microglia sculpt postnatal neural circuits in an activity and complement-dependent manner. Neuron, 74(4), 691–705. 10.1016/j.neuron.2012.03.026

Schneider, C. A., Rasband, W. S., & Eliceiri, K. W. (2012). NIH Image to ImageJ: 25 years of image analysis. Nature Methods, 9(7), Article 7. 10.1038/nmeth.2089

Schwarz, J. M., Sholar, P. W., & Bilbo, S. D. (2012). Sex differences in microglial colonization of the developing rat brain. Journal of Neurochemistry, 120(6), 948–963. 10.1111/j.1471-4159.2011.07630.x

Sierra, A., Encinas, J. M., Deudero, J. J. P., Chancey, J. H., Enikolopov, G., Overstreet-Wadiche, L. S., Tsirka, S. E., & Maletic-Savatic, M. (2010). Microglia shape adult hippocampal neurogenesis through apoptosis-coupled phagocytosis. Cell Stem Cell, 7(4), 483–495. 10.1016/j.stem.2010.08.014

Sullivan, O., & Ciernia, A. V. (2022). Work hard, play hard: How sexually differentiated microglia work to shape social play and reproductive behavior. Frontiers in Behavioral Neuroscience, 16, 989011. 10.3389/fnbeh.2022.989011

Suzuki, K., Sugihara, G., Ouchi, Y., Nakamura, K., Futatsubashi, M., Takebayashi, K., Yoshihara, Y., Omata, K., Matsumoto, K., Tsuchiya, K. J., Iwata, Y., Tsujii, M., Sugiyama, T., & Mori, N. (2013). Microglial activation in young adults with autism spectrum disorder. JAMA Psychiatry, 70(1), 49–58. 10.1001/jamapsychiatry.2013.272

Taylor, S. E., Morganti-Kossmann, C., Lifshitz, J., & Ziebell, J. M. (2014). Rod Microglia: A Morphological Definition. PLoS ONE, 9(5), e97096. 10.1371/journal.pone.0097096

Terstege, D. J., Oboh, D. O., & Epp, J. R. (2022). FASTMAP: Open-Source Flexible Atlas Segmentation Tool for Multi-Area Processing of Biological Images. eNeuro, 9(2), ENEURO.0325-21.2022. 10.1523/ENEURO.0325-21.2022

Tetreault, N. A., Hakeem, A. Y., Jiang, S., Williams, B. A., Allman, E., Wold, B. J., & Allman, J. M. (2012). Microglia in the cerebral cortex in autism. Journal of Autism and Developmental Disorders, 42(12), 2569–2584. 10.1007/s10803-012-1513-0

Torres-Platas, S. G., Comeau, S., Rachalski, A., Bo, G. D., Cruceanu, C., Turecki, G., Giros, B., & Mechawar, N. (2014). Morphometric characterization of microglial phenotypes in human cerebral cortex. Journal of Neuroinflammation, 11(1), 12. 10.1186/1742-2094-11-12

Wang, C., Yue, H., Hu, Z., Shen, Y., Ma, J., Li, J., Wang, X. D., Wang, L., Sun, B., Shi, P., Wang, L., & Gu, Y. (2020). Microglia mediate forgetting via complement-dependent synaptic elimination. Science, 367(6478), 688–694. 10.1126/science.aaz2288

Wendeln, A.-C., Degenhardt, K., Kaurani, L., Gertig, M., Ulas, T., Jain, G., Wagner, J., Häsler, L. M., Wild, K., Skodras, A., Blank, T., Staszewski, O., Datta, M., Centeno, T. P., Capece, V., Islam, M. R., Kerimoglu, C., Staufenbiel, M., Schultze, J. L., … Neher, J. J. (2018). Innate immune memory in the brain shapes neurological disease hallmarks. Nature, 556(7701), 332–338. 10.1038/s41586-018-0023-4

Wickham, H., Averick, M., Bryan, J., Chang, W., McGowan, L. D., François, R., Grolemund, G., Hayes, A., Henry, L., Hester, J., Kuhn, M., Pedersen, T. L., Miller, E., Bache, S. M., Müller, K., Ooms, J., Robinson, D., Seidel, D. P., Spinu, V., … Yutani, H. (2019). Welcome to the Tidyverse. Journal of Open Source Software, 4(43), 1686. 10.21105/joss.01686

York, E. M., LeDue, J. M., Bernier, L.-P., & MacVicar, B. A. (2018). 3DMorph Automatic Analysis of Microglial Morphology in Three Dimensions from Ex Vivo and In Vivo Imaging. eNeuro, 5(6), ENEURO.0266-18.2018. 10.1523/ENEURO.0266-18.2018

Young, K., & Morrison, H. (2018). Quantifying Microglia Morphology from Photomicrographs of Immunohistochemistry Prepared Tissue Using ImageJ. JoVE (Journal of Visualized Experiments), 136, e57648. 10.3791/57648

Zhan, Y., Paolicelli, R. C., Sforazzini, F., Weinhard, L., Bolasco, G., Pagani, F., Vyssotski, A. L., Bifone, A., Gozzi, A., Ragozzino, D., & Gross, C. T. (2014). Deficient neuron-microglia signaling results in impaired functional brain connectivity and social behavior. Nature Neuroscience, 17(3), 400–406. 10.1038/nn.3641

Zhuo, C., Tian, H., Song, X., Jiang, D., Chen, G., Cai, Z., Ping, J., Cheng, L., Zhou, C., & Chen, C. (2023). Microglia and cognitive impairment in schizophrenia: Translating scientific progress into novel therapeutic interventions. Schizophrenia, 9(1), Article 1. 10.1038/s41537-023-00370-z

