## Supplementary Info 1 for "Development of a high-throughput pipeline to characterize microglia morphological states at a single-cell resolution": SuppInfo1_Kim2023_ImageTypeComp.html

Kim2023\_ImageTypeComparisons


### Kim2023\_ImageTypeComparisons

#### 2023-09-24

##### Load libraries

```
library(tidyverse)
library(pheatmap)
library(factoextra)
library(nlme)
library(ggplot2)
library(hrbrthemes)
library(viridis)
library(ggforce)
library(lme4)
library(lmerTest)
library(Hmisc)
library(plotly)
library(ggpubr)
library(gridExtra)
library(MicrogliaMorphologyR)
```

##### Set seed for reproducibility

```
set.seed(1)
```

##### Load ImageTypeComparison dataset

```
data <- MicrogliaMorphologyR::data_ImageTypeComparison
```

##### PCA

```
data_log <- transform_log(data, x=1, start=5, end=13)
```

```
## Warning: `funs()` was deprecated in dplyr 0.8.0.
## ℹ Please use a list of either functions or lambdas:
## 
## # Simple named list: list(mean = mean, median = median)
## 
## # Auto named with `tibble::lst()`: tibble::lst(mean, median)
## 
## # Using lambdas list(~ mean(., trim = .2), ~ median(., na.rm = TRUE))
## ℹ The deprecated feature was likely used in the MicrogliaMorphologyR package.
##   Please report the issue to the authors.
## This warning is displayed once every 8 hours.
## Call `lifecycle::last_lifecycle_warnings()` to see where this warning was
## generated.
```

```
pcadata_elbow(data_log, featurestart=5, featureend=13)
```

```
pca_data <- pcadata(data_log, featurestart=5, featureend=13,
                    pc.start=1, pc.end=5)
pca_data %>%
  ggplot(aes(x=PC1, y=PC2, color=MorphologyClass)) +
  geom_point(size=2.5) +
  scale_colour_viridis_d() +
  #ggtitle("K-means clusters") +
  labs(color="Morphology Class") +
  theme_classic(base_size=16)
```

```
pca_data %>%
  ggplot(aes(x=PC1, y=PC2, color=ImageType)) +
  geom_point(size=2.5) +
  scale_colour_viridis_d() +
  #ggtitle("K-means clusters") +
  labs(color="Image Type") +
  theme_classic(base_size=16)
```

### PC by Feature correlations and Image type correlations

```
pcfeaturecorrelations(pca_data, pc.start=1, pc.end=5,
                      feature.start=10, feature.end=18,
                      rthresh=0.8, pthresh=0.05,
                      title="Spearman correlations")
```

```
## Correlation heatmap on PCs
edf <- pca_data %>% filter(ImageType=="EDF")
colnames(edf)<- paste0(colnames(edf), "_EDF")

twod <- pca_data %>% filter(ImageType=="2D")
colnames(twod)<- paste0(colnames(twod), "_2D")

threed <- pca_data %>% filter(ImageType=="3D")
colnames(threed)<- paste0(colnames(threed), "_3D")

input <- cbind(edf[,1:2], twod[,1:2], threed[,1:2])
input <- input[,order(colnames(input))]

hi <- rcorr(as.matrix(scale(input), type="pearson"))

mat1 <- hi$r
filter <- which(abs(mat1)<0.7)

mat2 <- hi$P
mat2 <- round(mat2,3)

mat2[mat2 < 0.05] <- "*" # significant p-values
mat2[mat2 > 0.05] <- "" # insignificant p-values
mat2[is.na(mat2)] <- "" # NAs (should be 27)
mat2[filter] <- "" # correlation values that are less than 0.5 (weak correlations)

# make heatmap
pheatmap(hi$r, display_numbers = mat2, fontsize_number=24,
         #cluster_rows=FALSE, cluster_cols=FALSE,
         fontsize=10, fontsize_row=10, fontsize_col=10,
         main="Pearson's correlation of PCs across Image Types")
```

### Boxplots and lineplots for differences across morphology subtypes

```
df2 <- data %>% group_by(ImageType) %>%
  mutate_at(5:13, scale)

data2 <- df2 %>% gather(measure, value, 5:13)
data2$ImageType <- factor(data2$ImageType, levels=c("3D","EDF","2D"))
data2$MorphologyClass <- factor(data2$MorphologyClass)

# all 3 image types (min-max normalized)
data2 %>%
  ggplot(aes(x=MorphologyClass, y=value, group=interaction(MorphologyClass,ImageType))) +
  facet_wrap(~measure, scales="free") +
  geom_boxplot(aes(group=interaction(MorphologyClass,ImageType), fill=ImageType), outlier.shape=NA) +
  scale_fill_viridis_d() +
  geom_point(position=position_dodge(width=0.8), size=0.75, aes(group=interaction(MorphologyClass,ImageType))) +
  ggtitle("Image type comparison") +
  labs(fill="Image Type") +
  theme_classic(base_size=14) +
  theme(axis.text.x=element_text(angle=45, vjust=1, hjust=1)) +
  theme(strip.background=element_rect(fill="#CCCCCC")) +
  ylab("Values scaled within image types")
```

```
# line graphs
df2 <- data %>% gather(measure, value, 5:13)
df2$ImageType <- factor(df2$ImageType, levels=c("3D","EDF","2D"))
df2$MorphologyClass <- factor(df2$MorphologyClass)

df2 %>% 
  ggplot(aes(x=ImageType, y=value, group=CellID, color=MorphologyClass)) +
  geom_line() +
  facet_wrap(~measure, scales="free") +
  scale_color_viridis_d() +
  ggtitle("Image type comparison") +
  labs(fill="Image Type") +
  theme_classic(base_size=14) +
  theme(axis.text.x=element_text(angle=45, vjust=1, hjust=1)) +
  theme(strip.background=element_rect(fill="#CCCCCC")) +
  ylab("Value")
```

##### Rcorr line plots

```
data2 <- data %>% gather(measure, value, 5:13)
data2$ImageType <- factor(data2$ImageType, levels=c("3D","EDF","2D"))
data2$MorphologyClass <- factor(data2$MorphologyClass)

## Correlation heatmap on numerical features 
edf <- data %>% filter(ImageType=="EDF")
colnames(edf)<- paste0(colnames(edf), "_EDF")

twod <- data %>% filter(ImageType=="2D")
colnames(twod)<- paste0(colnames(twod), "_2D")

threed <- data %>% filter(ImageType=="3D")
colnames(threed)<- paste0(colnames(threed), "_3D")

input <- cbind(edf[,5:13], twod[,5:13], threed[,5:13])
input <- input[,order(colnames(input))]

featurenames <- c("# of branches",
                   "# of junctions",
                   "# of end point voxels",
                   "# of junction voxels",
                   "# of slab voxels",
                   "Average branch length",
                   "# of triple points",
                   "# of quadruple points",
                   "Maximum branch length")

twoDthreeD = list()
twoDedf = list()
threeDedf = list()
for(f in featurenames){
  twoDthreeD[[f]] <-
    ggscatter(input, x = paste0(f,"_2D"), y = paste0(f,"_3D"),
            add = "reg.line", conf.int = TRUE,
            cor.coef = TRUE, cor.method = "pearson",
            title=f)
            
  twoDedf[[f]] <-
    ggscatter(input, x = paste0(f,"_2D"), y = paste0(f,"_EDF"),
            add = "reg.line", conf.int = TRUE,
            cor.coef = TRUE, cor.method = "pearson",
            title=f)
            
  threeDedf[[f]] <-
    ggscatter(input, x = paste0(f,"_3D"), y = paste0(f,"_EDF"),
            add = "reg.line", conf.int = TRUE,
            cor.coef = TRUE, cor.method = "pearson",
            title=f)
}

do.call("grid.arrange", c(twoDthreeD, ncol=3))
```

```
do.call("grid.arrange", c(twoDedf, ncol=3))
```

```
do.call("grid.arrange", c(threeDedf, ncol=3))
```
