## Supplementary Info 2 for "Development of a high-throughput pipeline to characterize microglia morphological states at a single-cell resolution": SuppInfo2_Kim2023_DataAnalysis_2xLPS.html


### Kim2023\_DataAnalysis\_2xLPS

###### Jenn Kim

#### 20/09/2023

##### load in libraries

```
library(MicrogliaMorphologyR)
library(factoextra)
library(DT)
```

### 2XLPS (0.5 mg/kg) DATA ANALYSIS

##### set seed

```
set.seed(1)
```

##### load in example dataset

```
data_2xLPS <- MicrogliaMorphologyR::data_2xLPS_mouse_fuzzykmeans
```

##### exploratory data visualization and data transformation for downstream analyses

```
# gather your numerical morphology data into one column ('measure') which contains the feature name, and another column ('value') which contains measured values
data_2xLPS_gathered <- data_2xLPS %>% gather(measure, value, 9:35)

# check for outliers and feature distributions
outliers_boxplots(data_2xLPS_gathered) # area and foreground pixels dominate analysis
```

```
outliers_distributions(data_2xLPS_gathered)
```

```
normalize_logplots(data_2xLPS_gathered,1) # after taking log(feature + 1)
```

```
# transform your data in appropriate manner for downstream analyses
data_2xLPS_logtransformed <- transform_log(data_2xLPS, 1, start=9, end=35) # we will use the logtransformed data as our PCA input

# get sample size of data based on factors of interest
samplesize(data_2xLPS, MouseID, Treatment, Antibody)
```

```
## # A tibble: 18 × 4
## # Groups:   MouseID, Treatment [6]
##    MouseID Treatment Antibody   num
##    <chr>   <chr>     <chr>    <int>
##  1 1       2xLPS     Cx3cr1    1703
##  2 1       2xLPS     Iba1      1737
##  3 1       2xLPS     P2ry12    2105
##  4 2       PBS       Cx3cr1    2496
##  5 2       PBS       Iba1      2927
##  6 2       PBS       P2ry12    4341
##  7 3       PBS       Cx3cr1    1145
##  8 3       PBS       Iba1      1310
##  9 3       PBS       P2ry12    1978
## 10 4       2xLPS     Cx3cr1    1775
## 11 4       2xLPS     Iba1      2044
## 12 4       2xLPS     P2ry12    2372
## 13 5       PBS       Cx3cr1    2053
## 14 5       PBS       Iba1      2302
## 15 5       PBS       P2ry12    3513
## 16 6       2xLPS     Cx3cr1    2771
## 17 6       2xLPS     Iba1      3095
## 18 6       2xLPS     P2ry12    3665
```

```
samplesize(data_2xLPS, Treatment, Antibody, BrainRegion, Subregion)
```

```
## # A tibble: 54 × 5
## # Groups:   Treatment, Antibody, BrainRegion [18]
##    Treatment Antibody BrainRegion Subregion   num
##    <chr>     <chr>    <chr>       <chr>     <int>
##  1 2xLPS     Cx3cr1   FC          ACC         305
##  2 2xLPS     Cx3cr1   FC          IL          517
##  3 2xLPS     Cx3cr1   FC          PL          405
##  4 2xLPS     Cx3cr1   HC          CA1         693
##  5 2xLPS     Cx3cr1   HC          CA2         175
##  6 2xLPS     Cx3cr1   HC          CA3         440
##  7 2xLPS     Cx3cr1   HC          DG          650
##  8 2xLPS     Cx3cr1   STR         CP         2052
##  9 2xLPS     Cx3cr1   STR         NA         1012
## 10 2xLPS     Iba1     FC          ACC         349
## # ℹ 44 more rows
```

##### generate heatmap of correlations across features

```
featurecorrelations(data_2xLPS, featurestart=9, featureend=35, rthresh=0.8, pthresh=0.05, title="Correlations across features")
```

##### variability described by PCs

```
pcadata_elbow(data_2xLPS_logtransformed, featurestart=9, featureend=35)
```

##### generate heatmap of correlations between PCs and features

```
pcfeaturecorrelations(data_2xLPS, pc.start=36, pc.end=38, 
                      feature.start=9, feature.end=35, 
                      rthresh=0.75, pthresh=0.05, 
                      title="Correlation between PCs and features")
```

##### visually explore different sources of variability in dataset

```
# gather your data by experimental variables (e.g., Treatment, Sex, MouseID, etc.)
gathered_expvariables <- data_2xLPS %>% gather(variable, value, 1:6) 

plots_expvariable(gathered_expvariables, "PC1", "PC2")
```

##### Cluster optimization

```
## for k-means clustering: scale PCs 1-3, which together describe ~85% of variability
pca_data_scale <- transform_scale(data_2xLPS, start=36, end=38) # scale pca data as input for k-means clustering
kmeans_input <- pca_data_scale[36:38]

# check for optimal number of clusters using wss and silhouette methods
sampling <- kmeans_input[sample(nrow(kmeans_input), 5000),] #sample 5000 random rows for cluster optimization

fviz_nbclust(sampling, kmeans, method = 'wss', nstart=25, iter.max=50) # 4 clusters
```

```
fviz_nbclust(sampling, kmeans, method = 'silhouette', nstart=25, iter.max=50) # 4 clusters
```

##### Plot k-means clusters in PC space

```
plot <- clusterplots(data_2xLPS, "PC1", "PC2")
plot + scale_colour_manual(values=c("#BBCC33","#44BB99","#EEDD88","#EE8866"))
```

##### Cluster-specific measures on average for each morphology feature, relative to other clusters

```
clusterfeatures(data_2xLPS, featurestart=9, featureend=35)
```

##### Cluster characterization and stats

### Brain regions

```
# calculate cluster percentages across variables of interest
cp <- clusterpercentage(data_2xLPS, "Cluster", MouseID, Antibody, Treatment, Sex, BrainRegion)
cp$Treatment <- factor(cp$Treatment, levels=c("PBS","2xLPS"))

# update cluster labels
cp <- cp %>% mutate(Cluster = 
                      case_when(Cluster=="1" ~ "Ameboid",
                                Cluster=="2" ~ "Rod-like",
                                Cluster=="3" ~ "Hypertrophic",
                                Cluster=="4" ~ "Ramified"))

cp$BrainRegion <- factor(cp$BrainRegion, levels=c("FC","HC","STR"))

# example graph of data given variables of interest
cp %>% 
  ggplot(aes(x=Cluster, y=percentage, group=interaction(Cluster, Treatment))) +
  facet_wrap(~Antibody*BrainRegion, ncol=3, scales="free") +
  geom_boxplot(aes(group=interaction(Cluster, Treatment), fill=Treatment)) +
  scale_fill_manual(values=c("#fde725","#482878")) +
  geom_point(position=position_dodge(width=0.8), size=0.75, aes(group=interaction(Cluster,Treatment), color=Sex)) +
  ggtitle("2xLPS mouse dataset: K-means clusters") +
  labs(fill="Treatment") +
  theme_classic(base_size=12) +
  theme(axis.text.x=element_text(angle=45, vjust=1, hjust=1)) +
  theme(strip.background=element_rect(fill="#CCCCCC")) +
  ylim(0.05,0.5)
```

#### Statistical analysis

##### Cluster percentage changes at animal level, in response to experimental variables

###### e.g., Across clusters - How does cluster membership change with LPS?

```
# prepare percentages dataset for downstream analysis
stats.input <- cp
stats.input$MouseID <- factor(stats.input$MouseID)
stats.input$Cluster <- factor(stats.input$Cluster)
stats.input$Treatment <- factor(stats.input$Treatment)
stats.input$BrainRegion <- factor(stats.input$BrainRegion)
stats.input$Antibody <- factor(stats.input$Antibody)

# run stats analysis for changes in cluster percentages, at the animal level
# you can specify up to two posthoc comparisons (posthoc1 and posthoc2 arguments) - if you only have one set of posthocs to run, specify the same comparison twice for both arguments. you will just get the same results in output[[2]] and output[[3]].
stats.testing <- stats_cluster.animal(stats.input, "percentage ~ Cluster*Treatment*BrainRegion + Antibody + (1|MouseID)", 
                                      "~Cluster*Treatment*BrainRegion", "~Cluster*Treatment*BrainRegion", "bonferroni")
```

```
## Formula:          
## percentage ~ Cluster * Treatment * BrainRegion + Antibody + (1 |      MouseID)
## Data: data
##       AIC       BIC    logLik  df.resid 
## -569.0465 -476.1392  312.5233       176 
## Random-effects (co)variances:
## 
## Conditional model:
##  Groups  Name        Std.Dev. 
##  MouseID (Intercept) 4.551e-06
## 
## Number of obs: 204 / Conditional model: MouseID, 6
## 
## Dispersion parameter for beta family (): 62.4 
## 
## Fixed Effects:
## 
## Conditional model:
##                      (Intercept)                          Cluster1  
##                       -1.1449978                        -0.1984451  
##                         Cluster2                          Cluster3  
##                       -0.2334864                         0.3788346  
##                       Treatment1                      BrainRegion1  
##                       -0.0146812                         0.0087407  
##                     BrainRegion2                         Antibody1  
##                        0.0090323                         0.0008063  
##                        Antibody2               Cluster1:Treatment1  
##                        0.0069276                         0.3449797  
##              Cluster2:Treatment1               Cluster3:Treatment1  
##                       -0.5071318                         0.0557719  
##            Cluster1:BrainRegion1             Cluster2:BrainRegion1  
##                       -0.0163807                         0.0549738  
##            Cluster3:BrainRegion1             Cluster1:BrainRegion2  
##                       -0.0950720                        -0.0392781  
##            Cluster2:BrainRegion2             Cluster3:BrainRegion2  
##                        0.2404445                        -0.1916492  
##          Treatment1:BrainRegion1           Treatment1:BrainRegion2  
##                       -0.0025026                         0.0071379  
## Cluster1:Treatment1:BrainRegion1  Cluster2:Treatment1:BrainRegion1  
##                       -0.0062024                        -0.0261938  
## Cluster3:Treatment1:BrainRegion1  Cluster1:Treatment1:BrainRegion2  
##                        0.1319542                        -0.0187652  
## Cluster2:Treatment1:BrainRegion2  Cluster3:Treatment1:BrainRegion2  
##                       -0.0844890                         0.0598352
```

```
stats.testing[[1]] # anova
```

```
## Analysis of Deviance Table (Type II Wald chisquare tests)
## 
## Response: percentage
##                                  Chisq Df Pr(>Chisq)    
## Cluster                       108.4426  3  < 2.2e-16 ***
## Treatment                       0.0005  1   0.982901    
## BrainRegion                     0.0376  2   0.981384    
## Antibody                        0.0845  2   0.958629    
## Cluster:Treatment             191.0931  3  < 2.2e-16 ***
## Cluster:BrainRegion            64.8611  6  4.605e-12 ***
## Treatment:BrainRegion           0.6413  2   0.725672    
## Cluster:Treatment:BrainRegion  20.4785  6   0.002275 ** 
## ---
## Signif. codes:  0 '***' 0.001 '**' 0.01 '*' 0.05 '.' 0.1 ' ' 1
```

```
stats.testing[[5]] # summary of model
```

```
## Formula:          
## percentage ~ Cluster * Treatment * BrainRegion + Antibody + (1 |      MouseID)
## Data: data
##       AIC       BIC    logLik  df.resid 
## -569.0465 -476.1392  312.5233       176 
## Random-effects (co)variances:
## 
## Conditional model:
##  Groups  Name        Std.Dev. 
##  MouseID (Intercept) 4.551e-06
## 
## Number of obs: 204 / Conditional model: MouseID, 6
## 
## Dispersion parameter for beta family (): 62.4 
## 
## Fixed Effects:
## 
## Conditional model:
##                      (Intercept)                          Cluster1  
##                       -1.1449978                        -0.1984451  
##                         Cluster2                          Cluster3  
##                       -0.2334864                         0.3788346  
##                       Treatment1                      BrainRegion1  
##                       -0.0146812                         0.0087407  
##                     BrainRegion2                         Antibody1  
##                        0.0090323                         0.0008063  
##                        Antibody2               Cluster1:Treatment1  
##                        0.0069276                         0.3449797  
##              Cluster2:Treatment1               Cluster3:Treatment1  
##                       -0.5071318                         0.0557719  
##            Cluster1:BrainRegion1             Cluster2:BrainRegion1  
##                       -0.0163807                         0.0549738  
##            Cluster3:BrainRegion1             Cluster1:BrainRegion2  
##                       -0.0950720                        -0.0392781  
##            Cluster2:BrainRegion2             Cluster3:BrainRegion2  
##                        0.2404445                        -0.1916492  
##          Treatment1:BrainRegion1           Treatment1:BrainRegion2  
##                       -0.0025026                         0.0071379  
## Cluster1:Treatment1:BrainRegion1  Cluster2:Treatment1:BrainRegion1  
##                       -0.0062024                        -0.0261938  
## Cluster3:Treatment1:BrainRegion1  Cluster1:Treatment1:BrainRegion2  
##                        0.1319542                        -0.0187652  
## Cluster2:Treatment1:BrainRegion2  Cluster3:Treatment1:BrainRegion2  
##                       -0.0844890                         0.0598352
```

```
##### FOR EACH ANTIBODY #####
stats.input <- cp
stats.input$MouseID <- factor(stats.input$MouseID)
stats.input$Cluster <- factor(stats.input$Cluster)
stats.input$Treatment <- factor(stats.input$Treatment)
stats.input$BrainRegion <- factor(stats.input$BrainRegion)

# Iba1
stats.input2 <- stats.input %>% filter(Antibody=="Iba1")
stats.testing <- stats_cluster.animal(stats.input2, "percentage ~ Cluster*Treatment*BrainRegion + (1|MouseID)", 
                                      "~Treatment|Cluster|BrainRegion", "~Treatment|Cluster|BrainRegion", "bonferroni")
```

```
## Formula:          
## percentage ~ Cluster * Treatment * BrainRegion + (1 | MouseID)
## Data: data
##       AIC       BIC    logLik  df.resid 
## -245.5320 -187.8248  148.7660        42 
## Random-effects (co)variances:
## 
## Conditional model:
##  Groups  Name        Std.Dev. 
##  MouseID (Intercept) 2.857e-06
## 
## Number of obs: 68 / Conditional model: MouseID, 6
## 
## Dispersion parameter for beta family ():  237 
## 
## Fixed Effects:
## 
## Conditional model:
##                      (Intercept)                          Cluster1  
##                        -1.145548                         -0.397575  
##                         Cluster2                          Cluster3  
##                        -0.040224                          0.312574  
##                       Treatment1                      BrainRegion1  
##                         0.007090                         -0.009871  
##                     BrainRegion2               Cluster1:Treatment1  
##                         0.010161                          0.240106  
##              Cluster2:Treatment1               Cluster3:Treatment1  
##                        -0.496992                          0.122607  
##            Cluster1:BrainRegion1             Cluster2:BrainRegion1  
##                         0.058172                         -0.069723  
##            Cluster3:BrainRegion1             Cluster1:BrainRegion2  
##                        -0.101565                         -0.064662  
##            Cluster2:BrainRegion2             Cluster3:BrainRegion2  
##                         0.296681                         -0.174165  
##          Treatment1:BrainRegion1           Treatment1:BrainRegion2  
##                        -0.013858                          0.015080  
## Cluster1:Treatment1:BrainRegion1  Cluster2:Treatment1:BrainRegion1  
##                         0.053383                         -0.128580  
## Cluster3:Treatment1:BrainRegion1  Cluster1:Treatment1:BrainRegion2  
##                         0.173579                         -0.066085  
## Cluster2:Treatment1:BrainRegion2  Cluster3:Treatment1:BrainRegion2  
##                        -0.018685                          0.031872
```

```
stats.testing[[2]]
```

```
##  contrast    Cluster      BrainRegion   estimate        SE  df z.ratio p.value
##  PBS - 2xLPS Ameboid      FC           0.5734414 0.1382056 Inf   4.149  0.0004
##  PBS - 2xLPS Hypertrophic FC          -1.2646789 0.1355486 Inf  -9.330  <.0001
##  PBS - 2xLPS Ramified     FC           0.5788350 0.1190877 Inf   4.861  <.0001
##  PBS - 2xLPS Rod-like     FC           0.0582594 0.1169659 Inf   0.498  1.0000
##  PBS - 2xLPS Ameboid      HC           0.3923815 0.1565036 Inf   2.507  0.1460
##  PBS - 2xLPS Hypertrophic HC          -0.9870125 0.1368854 Inf  -7.211  <.0001
##  PBS - 2xLPS Ramified     HC           0.3532981 0.1326180 Inf   2.664  0.0927
##  PBS - 2xLPS Rod-like     HC           0.4186942 0.1348397 Inf   3.105  0.0228
##  PBS - 2xLPS Ameboid      STR          0.5173534 0.1397094 Inf   3.703  0.0026
##  PBS - 2xLPS Hypertrophic STR         -0.6877177 0.1355466 Inf  -5.074  <.0001
##  PBS - 2xLPS Ramified     STR         -0.1539517 0.1099344 Inf  -1.400  1.0000
##  PBS - 2xLPS Rod-like     STR          0.3712634 0.1218959 Inf   3.046  0.0279
##  Significant
##  significant
##  significant
##  significant
##  ns         
##  ns         
##  significant
##  ns         
##  significant
##  significant
##  significant
##  ns         
##  significant
## 
## Results are given on the log odds ratio (not the response) scale. 
## P value adjustment: bonferroni method for 12 tests
```

```
# Cx3cr1
stats.input2 <- stats.input %>% filter(Antibody=="Cx3cr1")
stats.testing <- stats_cluster.animal(stats.input2, "percentage ~ Cluster*Treatment*BrainRegion + (1|MouseID)", 
                                      "~Treatment|Cluster|BrainRegion", "~Treatment|Cluster|BrainRegion", "bonferroni")
```

```
## Formula:          
## percentage ~ Cluster * Treatment * BrainRegion + (1 | MouseID)
## Data: data
##       AIC       BIC    logLik  df.resid 
## -273.8875 -216.1803  162.9437        42 
## Random-effects (co)variances:
## 
## Conditional model:
##  Groups  Name        Std.Dev. 
##  MouseID (Intercept) 2.978e-06
## 
## Number of obs: 68 / Conditional model: MouseID, 6
## 
## Dispersion parameter for beta family ():  343 
## 
## Fixed Effects:
## 
## Conditional model:
##                      (Intercept)                          Cluster1  
##                        -1.190055                         -0.495287  
##                         Cluster2                          Cluster3  
##                        -0.390526                          0.700924  
##                       Treatment1                      BrainRegion1  
##                        -0.022015                          0.024557  
##                     BrainRegion2               Cluster1:Treatment1  
##                         0.010409                          0.258197  
##              Cluster2:Treatment1               Cluster3:Treatment1  
##                        -0.546997                          0.154786  
##            Cluster1:BrainRegion1             Cluster2:BrainRegion1  
##                        -0.046294                          0.224384  
##            Cluster3:BrainRegion1             Cluster1:BrainRegion2  
##                        -0.163178                         -0.008548  
##            Cluster2:BrainRegion2             Cluster3:BrainRegion2  
##                         0.125492                         -0.173133  
##          Treatment1:BrainRegion1           Treatment1:BrainRegion2  
##                         0.006138                         -0.007396  
## Cluster1:Treatment1:BrainRegion1  Cluster2:Treatment1:BrainRegion1  
##                        -0.035714                          0.028902  
## Cluster3:Treatment1:BrainRegion1  Cluster1:Treatment1:BrainRegion2  
##                         0.093719                          0.003264  
## Cluster2:Treatment1:BrainRegion2  Cluster3:Treatment1:BrainRegion2  
##                        -0.141311                          0.127014
```

```
stats.testing[[2]]
```

```
##  contrast    Cluster      BrainRegion   estimate         SE  df z.ratio p.value
##  PBS - 2xLPS Ameboid      FC           0.4132111 0.12274058 Inf   3.367  0.0091
##  PBS - 2xLPS Hypertrophic FC          -1.0679430 0.11321842 Inf  -9.433  <.0001
##  PBS - 2xLPS Ramified     FC           0.4652548 0.09308351 Inf   4.998  <.0001
##  PBS - 2xLPS Rod-like     FC           0.0624617 0.09913239 Inf   0.630  1.0000
##  PBS - 2xLPS Ameboid      HC           0.4640978 0.13424936 Inf   3.457  0.0066
##  PBS - 2xLPS Hypertrophic HC          -1.4354370 0.14131267 Inf -10.158  <.0001
##  PBS - 2xLPS Ramified     HC           0.5047767 0.10377351 Inf   4.864  <.0001
##  PBS - 2xLPS Rod-like     HC           0.2312729 0.10908371 Inf   2.120  0.4079
##  PBS - 2xLPS Ameboid      STR          0.5397795 0.12165487 Inf   4.437  0.0001
##  PBS - 2xLPS Hypertrophic STR         -0.9106913 0.13883712 Inf  -6.559  <.0001
##  PBS - 2xLPS Ramified     STR         -0.1734086 0.08852922 Inf  -1.959  0.6017
##  PBS - 2xLPS Rod-like     STR          0.3782609 0.10170895 Inf   3.719  0.0024
##  Significant
##  significant
##  significant
##  significant
##  ns         
##  significant
##  significant
##  significant
##  ns         
##  significant
##  significant
##  ns         
##  significant
## 
## Results are given on the log odds ratio (not the response) scale. 
## P value adjustment: bonferroni method for 12 tests
```

```
# P2ry12
stats.input2 <- stats.input %>% filter(Antibody=="P2ry12")
stats.testing <- stats_cluster.animal(stats.input2, "percentage ~ Cluster*Treatment*BrainRegion + (1|MouseID)", 
                                      "~Treatment|Cluster|BrainRegion", "~Treatment|Cluster|BrainRegion", "bonferroni")
```

```
## Formula:          
## percentage ~ Cluster * Treatment * BrainRegion + (1 | MouseID)
## Data: data
##       AIC       BIC    logLik  df.resid 
## -250.8298 -193.1226  151.4149        42 
## Random-effects (co)variances:
## 
## Conditional model:
##  Groups  Name        Std.Dev. 
##  MouseID (Intercept) 2.744e-06
## 
## Number of obs: 68 / Conditional model: MouseID, 6
## 
## Dispersion parameter for beta family ():  253 
## 
## Fixed Effects:
## 
## Conditional model:
##                      (Intercept)                          Cluster1  
##                       -1.157e+00                         2.883e-01  
##                         Cluster2                          Cluster3  
##                       -2.939e-01                         1.489e-01  
##                       Treatment1                      BrainRegion1  
##                       -3.129e-02                         1.317e-02  
##                     BrainRegion2               Cluster1:Treatment1  
##                        8.750e-03                         5.658e-01  
##              Cluster2:Treatment1               Cluster3:Treatment1  
##                       -5.188e-01                        -1.056e-01  
##            Cluster1:BrainRegion1             Cluster2:BrainRegion1  
##                       -6.356e-02                         1.558e-02  
##            Cluster3:BrainRegion1             Cluster1:BrainRegion2  
##                       -2.585e-02                        -4.892e-02  
##            Cluster2:BrainRegion2             Cluster3:BrainRegion2  
##                        3.167e-01                        -2.388e-01  
##          Treatment1:BrainRegion1           Treatment1:BrainRegion2  
##                       -8.361e-05                         1.591e-02  
## Cluster1:Treatment1:BrainRegion1  Cluster2:Treatment1:BrainRegion1  
##                       -3.684e-02                         2.176e-02  
## Cluster3:Treatment1:BrainRegion1  Cluster1:Treatment1:BrainRegion2  
##                        1.347e-01                         4.621e-03  
## Cluster2:Treatment1:BrainRegion2  Cluster3:Treatment1:BrainRegion2  
##                       -9.348e-02                         2.179e-02
```

```
stats.testing[[2]]
```

```
##  contrast    Cluster      BrainRegion   estimate        SE  df z.ratio p.value
##  PBS - 2xLPS Ameboid      FC           0.9951388 0.1174172 Inf   8.475  <.0001
##  PBS - 2xLPS Hypertrophic FC          -1.0568501 0.1351403 Inf  -7.820  <.0001
##  PBS - 2xLPS Ramified     FC          -0.0045921 0.1159168 Inf  -0.040  1.0000
##  PBS - 2xLPS Rod-like     FC          -0.1846960 0.1218126 Inf  -1.516  1.0000
##  PBS - 2xLPS Ameboid      HC           1.1100402 0.1290878 Inf   8.599  <.0001
##  PBS - 2xLPS Hypertrophic HC          -1.2553468 0.1451939 Inf  -8.646  <.0001
##  PBS - 2xLPS Ramified     HC          -0.1983860 0.1378762 Inf  -1.439  1.0000
##  PBS - 2xLPS Rod-like     HC           0.2206439 0.1394987 Inf   1.582  1.0000
##  PBS - 2xLPS Ameboid      STR          1.1017554 0.1149003 Inf   9.589  <.0001
##  PBS - 2xLPS Hypertrophic STR         -0.9884376 0.1529288 Inf  -6.463  <.0001
##  PBS - 2xLPS Ramified     STR         -0.6183581 0.1114082 Inf  -5.550  <.0001
##  PBS - 2xLPS Rod-like     STR          0.1280967 0.1271023 Inf   1.008  1.0000
##  Significant
##  significant
##  significant
##  ns         
##  ns         
##  significant
##  significant
##  ns         
##  ns         
##  significant
##  significant
##  significant
##  ns         
## 
## Results are given on the log odds ratio (not the response) scale. 
## P value adjustment: bonferroni method for 12 tests
```

### Subregions

```
# calculate cluster percentages across variables of interest
cp <- clusterpercentage(data_2xLPS, "Cluster", MouseID, Antibody, Treatment, Sex, Subregion)
cp$Treatment <- factor(cp$Treatment, levels=c("PBS","2xLPS"))

# update cluster labels
cp <- cp %>% mutate(Cluster = 
                      case_when(Cluster=="1" ~ "Ameboid",
                                Cluster=="2" ~ "Rod-like",
                                Cluster=="3" ~ "Hypertrophic",
                                Cluster=="4" ~ "Ramified"))

cp$Subregion <- factor(cp$Subregion, levels=c("IL","PL","ACC","CP","NA","CA1","CA2","CA3","DG"))

# example graph of data given variables of interest
cp %>% 
  #filter(Antibody=="Iba1") %>%
  ggplot(aes(x=Cluster, y=percentage, group=interaction(Cluster, Treatment))) +
  facet_wrap(~Antibody*Subregion, ncol=9, scales="free") +
  geom_boxplot(aes(group=interaction(Cluster, Treatment), fill=Treatment)) +
  scale_fill_manual(values=c("#fde725","#482878")) +
  geom_point(position=position_dodge(width=0.8), size=0.75, aes(group=interaction(Cluster,Treatment), color=Sex)) +
  ggtitle("2xLPS mouse dataset: K-means clusters") +
  labs(fill="Treatment") +
  theme_classic(base_size=12) +
  theme(axis.text.x=element_text(angle=45, vjust=1, hjust=1)) +
  theme(strip.background=element_rect(fill="#CCCCCC")) +
  ylim(0.02,0.51)
```

### Individual morphology measures by Brain Region

```
# prepare data for downstream analysis
data <- data_2xLPS %>% 
  group_by(MouseID, Sex, Treatment, BrainRegion, Antibody) %>% 
  summarise(across("Foreground pixels":"Maximum branch length", ~mean(.x))) %>% 
  gather(Measure, Value, "Foreground pixels":"Maximum branch length")

# filter out data you want to run stats on and make sure to make any variables included in model into factors
stats.input <- data 
stats.input$Treatment <- factor(stats.input$Treatment, levels=c("PBS","2xLPS"))
stats.input$BrainRegion <- factor(stats.input$BrainRegion)
stats.input$Antibody <- factor(stats.input$Antibody)

# # of junctions
stats.input %>% filter(Measure=="# of junctions") %>%
  ggplot(aes(x=Treatment, y=Value, group=Treatment)) +
  facet_wrap(~Measure*Antibody*BrainRegion, scales="free") +
  geom_boxplot(aes(group=Treatment, fill=Treatment)) +
  scale_fill_manual(values=c("#fde725","#482878")) +
  geom_point(position=position_dodge(width=0.8), size=0.75, aes(group=Treatment, color=Sex)) +
  labs(fill="Treatment") +
  theme_classic(base_size=12) +
  theme(axis.text.x=element_text(angle=45, vjust=1, hjust=1)) +
  theme(strip.background=element_rect(fill="#CCCCCC")) + 
  ylim(10,21)
```

```
# Area
stats.input %>% filter(Measure=="Area") %>%
  ggplot(aes(x=Treatment, y=Value, group=Treatment)) +
  facet_wrap(~Measure*Antibody*BrainRegion, scales="free") +
  geom_boxplot(aes(group=Treatment, fill=Treatment)) +
  scale_fill_manual(values=c("#fde725","#482878")) +
  geom_point(position=position_dodge(width=0.8), size=0.75, aes(group=Treatment, color=Sex)) +
  labs(fill="Treatment") +
  theme_classic(base_size=12) +
  theme(axis.text.x=element_text(angle=45, vjust=1, hjust=1)) +
  theme(strip.background=element_rect(fill="#CCCCCC")) + 
  ylim(6605,11476)
```

```
# Circularity
stats.input %>% filter(Measure=="Circularity") %>%
  ggplot(aes(x=Treatment, y=Value, group=Treatment)) +
  facet_wrap(~Measure*Antibody*BrainRegion, scales="free") +
  geom_boxplot(aes(group=Treatment, fill=Treatment)) +
  scale_fill_manual(values=c("#fde725","#482878")) +
  geom_point(position=position_dodge(width=0.8), size=0.75, aes(group=Treatment, color=Sex)) +
  labs(fill="Treatment") +
  theme_classic(base_size=12) +
  theme(axis.text.x=element_text(angle=45, vjust=1, hjust=1)) +
  theme(strip.background=element_rect(fill="#CCCCCC")) + 
  ylim(0.75,0.82)
```

##### Individual morphology measures, at the animal level (averaged for each measure)

###### e.g., How does each individual morphology measure change with LPS treatment?

```
# prepare data for downstream analysis
data <- data_2xLPS %>% 
  group_by(MouseID, Sex, Treatment, BrainRegion, Antibody) %>% 
  summarise(across("Foreground pixels":"Maximum branch length", ~mean(.x))) %>% 
  gather(Measure, Value, "Foreground pixels":"Maximum branch length")

# filter out data you want to run stats on and make sure to make any variables included in model into factors
stats.input <- data 
stats.input$Treatment <- factor(stats.input$Treatment, levels=c("PBS","2xLPS"))
stats.input$BrainRegion <- factor(stats.input$BrainRegion)
stats.input$Antibody <- factor(stats.input$Antibody)

# run stats analysis for changes in individual morphology measures
# you can specify up to two posthoc comparisons (posthoc1 and posthoc2 arguments) - if you only have one set of posthocs to run, specify the same comparison twice for both arguments. you will just get the same results in output[[2]] and output[[3]].
stats.testing <- stats_morphologymeasures.animal(stats.input, "Value ~ Treatment*BrainRegion + Antibody", 
                                                 "~Treatment*BrainRegion", "~Treatment*BrainRegion", "bonferroni")
```

```
## [1] "Foreground pixels"
## <simpleError in leveneTest.formula(formula, data, center = center): Model must be completely crossed formula only.>
## [1] "Density of foreground pixels in hull area"
## <simpleError in leveneTest.formula(formula, data, center = center): Model must be completely crossed formula only.>
## [1] "Span ratio of hull (major/minor axis)"
## <simpleError in leveneTest.formula(formula, data, center = center): Model must be completely crossed formula only.>
## [1] "Maximum span across hull"
## <simpleError in leveneTest.formula(formula, data, center = center): Model must be completely crossed formula only.>
## [1] "Area"
## <simpleError in leveneTest.formula(formula, data, center = center): Model must be completely crossed formula only.>
## [1] "Perimeter"
## <simpleError in leveneTest.formula(formula, data, center = center): Model must be completely crossed formula only.>
## [1] "Circularity"
## <simpleError in leveneTest.formula(formula, data, center = center): Model must be completely crossed formula only.>
## [1] "Width of bounding rectangle"
## <simpleError in leveneTest.formula(formula, data, center = center): Model must be completely crossed formula only.>
## [1] "Height of bounding rectangle"
## <simpleError in leveneTest.formula(formula, data, center = center): Model must be completely crossed formula only.>
## [1] "Maximum radius from hull's center of mass"
## <simpleError in leveneTest.formula(formula, data, center = center): Model must be completely crossed formula only.>
## [1] "Max/min radii from hull's center of mass"
## <simpleError in leveneTest.formula(formula, data, center = center): Model must be completely crossed formula only.>
## [1] "Relative variation (CV) in radii from hull's center of mass"
## <simpleError in leveneTest.formula(formula, data, center = center): Model must be completely crossed formula only.>
## [1] "Mean radius"
## <simpleError in leveneTest.formula(formula, data, center = center): Model must be completely crossed formula only.>
## [1] "Diameter of bounding circle"
## <simpleError in leveneTest.formula(formula, data, center = center): Model must be completely crossed formula only.>
## [1] "Maximum radius from circle's center of mass"
## <simpleError in leveneTest.formula(formula, data, center = center): Model must be completely crossed formula only.>
## [1] "Max/min radii from circle's center of mass"
## <simpleError in leveneTest.formula(formula, data, center = center): Model must be completely crossed formula only.>
## [1] "Relative variation (CV) in radii from circle's center of mass"
## <simpleError in leveneTest.formula(formula, data, center = center): Model must be completely crossed formula only.>
## [1] "Mean radius from circle's center of mass"
## <simpleError in leveneTest.formula(formula, data, center = center): Model must be completely crossed formula only.>
## [1] "# of branches"
## <simpleError in leveneTest.formula(formula, data, center = center): Model must be completely crossed formula only.>
## [1] "# of junctions"
## <simpleError in leveneTest.formula(formula, data, center = center): Model must be completely crossed formula only.>
## [1] "# of end point voxels"
## <simpleError in leveneTest.formula(formula, data, center = center): Model must be completely crossed formula only.>
## [1] "# of junction voxels"
## <simpleError in leveneTest.formula(formula, data, center = center): Model must be completely crossed formula only.>
## [1] "# of slab voxels"
## <simpleError in leveneTest.formula(formula, data, center = center): Model must be completely crossed formula only.>
## [1] "Average branch length"
## <simpleError in leveneTest.formula(formula, data, center = center): Model must be completely crossed formula only.>
## [1] "# of triple points"
## <simpleError in leveneTest.formula(formula, data, center = center): Model must be completely crossed formula only.>
## [1] "# of quadruple points"
## <simpleError in leveneTest.formula(formula, data, center = center): Model must be completely crossed formula only.>
## [1] "Maximum branch length"
## <simpleError in leveneTest.formula(formula, data, center = center): Model must be completely crossed formula only.>
## 
## Call:
## lm(formula = as.formula(paste(y.model)), data = tmp)
## 
## Coefficients:
##             (Intercept)               Treatment1             BrainRegion1  
##                18.32208                 -1.04707                 -0.03196  
##            BrainRegion2                Antibody1                Antibody2  
##                 0.49244                  0.12384                  0.51207  
## Treatment1:BrainRegion1  Treatment1:BrainRegion2  
##                -0.07882                 -0.11006
```

```
stats.testing[[1]] %>% DT::datatable(., options=list(autoWidth=TRUE, scrollX=TRUE, scrollCollapse=TRUE)) # anova
```

```
stats.testing[[7]] # summary of model
```

```
## 
## Call:
## lm(formula = as.formula(paste(y.model)), data = tmp)
## 
## Coefficients:
##             (Intercept)               Treatment1             BrainRegion1  
##                18.32208                 -1.04707                 -0.03196  
##            BrainRegion2                Antibody1                Antibody2  
##                 0.49244                  0.12384                  0.51207  
## Treatment1:BrainRegion1  Treatment1:BrainRegion2  
##                -0.07882                 -0.11006
```

```
##### FOR EACH ANTIBODY #####
stats.input <- data
stats.input$Treatment <- factor(stats.input$Treatment, levels=c("PBS","2xLPS"))
stats.input$BrainRegion <- factor(stats.input$BrainRegion)

# Iba1
stats.input2 <- stats.input %>% filter(Antibody=="Iba1")
stats.testing <- stats_morphologymeasures.animal(stats.input2, "Value ~ Treatment*BrainRegion", 
                                      "~Treatment|BrainRegion", "~Treatment|BrainRegion", "bonferroni")
```

```
## [1] "Foreground pixels"
## [1] "Density of foreground pixels in hull area"
## [1] "Span ratio of hull (major/minor axis)"
## [1] "Maximum span across hull"
## [1] "Area"
## [1] "Perimeter"
## [1] "Circularity"
## [1] "Width of bounding rectangle"
## [1] "Height of bounding rectangle"
## [1] "Maximum radius from hull's center of mass"
## [1] "Max/min radii from hull's center of mass"
## [1] "Relative variation (CV) in radii from hull's center of mass"
## [1] "Mean radius"
## [1] "Diameter of bounding circle"
## [1] "Maximum radius from circle's center of mass"
## [1] "Max/min radii from circle's center of mass"
## [1] "Relative variation (CV) in radii from circle's center of mass"
## [1] "Mean radius from circle's center of mass"
## [1] "# of branches"
## [1] "# of junctions"
## [1] "# of end point voxels"
## [1] "# of junction voxels"
## [1] "# of slab voxels"
## [1] "Average branch length"
## [1] "# of triple points"
## [1] "# of quadruple points"
## [1] "Maximum branch length"
## 
## Call:
## lm(formula = as.formula(paste(y.model)), data = tmp)
## 
## Coefficients:
##             (Intercept)               Treatment1             BrainRegion1  
##                 18.8678                  -0.6961                  -0.2791  
##            BrainRegion2  Treatment1:BrainRegion1  Treatment1:BrainRegion2  
##                  0.6411                  -0.1174                  -0.0377
```

```
stats.testing[[2]] %>% DT::datatable(., options=list(autoWidth=TRUE, scrollX=TRUE, scrollCollapse=TRUE))
```

```
do.call("grid.arrange", c(stats.testing[[4]], ncol=4)) # qqplots to check normality assumptions
```

```
stats.testing[[5]] %>% DT::datatable(., options=list(autoWidth=TRUE, scrollX=TRUE, scrollCollapse=TRUE)) # levene
```

```
stats.testing[[6]] %>% DT::datatable(., options=list(autoWidth=TRUE, scrollX=TRUE, scrollCollapse=TRUE)) # shapiro
```

```
stats.testing[[7]] # summary of model
```

```
## 
## Call:
## lm(formula = as.formula(paste(y.model)), data = tmp)
## 
## Coefficients:
##             (Intercept)               Treatment1             BrainRegion1  
##                 18.8678                  -0.6961                  -0.2791  
##            BrainRegion2  Treatment1:BrainRegion1  Treatment1:BrainRegion2  
##                  0.6411                  -0.1174                  -0.0377
```

```
# Cx3cr1
stats.input2 <- stats.input %>% filter(Antibody=="Cx3cr1")
stats.testing <- stats_morphologymeasures.animal(stats.input2, "Value ~ Treatment*BrainRegion", 
                                      "~Treatment|BrainRegion", "~Treatment|BrainRegion", "bonferroni")
```

```
## [1] "Foreground pixels"
## [1] "Density of foreground pixels in hull area"
## [1] "Span ratio of hull (major/minor axis)"
## [1] "Maximum span across hull"
## [1] "Area"
## [1] "Perimeter"
## [1] "Circularity"
## [1] "Width of bounding rectangle"
## [1] "Height of bounding rectangle"
## [1] "Maximum radius from hull's center of mass"
## [1] "Max/min radii from hull's center of mass"
## [1] "Relative variation (CV) in radii from hull's center of mass"
## [1] "Mean radius"
## [1] "Diameter of bounding circle"
## [1] "Maximum radius from circle's center of mass"
## [1] "Max/min radii from circle's center of mass"
## [1] "Relative variation (CV) in radii from circle's center of mass"
## [1] "Mean radius from circle's center of mass"
## [1] "# of branches"
## [1] "# of junctions"
## [1] "# of end point voxels"
## [1] "# of junction voxels"
## [1] "# of slab voxels"
## [1] "Average branch length"
## [1] "# of triple points"
## [1] "# of quadruple points"
## [1] "Maximum branch length"
## 
## Call:
## lm(formula = as.formula(paste(y.model)), data = tmp)
## 
## Coefficients:
##             (Intercept)               Treatment1             BrainRegion1  
##                18.43305                 -0.96622                  0.13381  
##            BrainRegion2  Treatment1:BrainRegion1  Treatment1:BrainRegion2  
##                 0.31104                 -0.03636                 -0.22824
```

```
stats.testing[[2]] %>% DT::datatable(., options=list(autoWidth=TRUE, scrollX=TRUE, scrollCollapse=TRUE))
```

```
do.call("grid.arrange", c(stats.testing[[4]], ncol=4)) # qqplots to check normality assumptions
```

```
stats.testing[[5]] %>% DT::datatable(., options=list(autoWidth=TRUE, scrollX=TRUE, scrollCollapse=TRUE)) # levene
```

```
stats.testing[[6]] %>% DT::datatable(., options=list(autoWidth=TRUE, scrollX=TRUE, scrollCollapse=TRUE)) # shapiro
```

```
stats.testing[[7]] # summary of model
```

```
## 
## Call:
## lm(formula = as.formula(paste(y.model)), data = tmp)
## 
## Coefficients:
##             (Intercept)               Treatment1             BrainRegion1  
##                18.43305                 -0.96622                  0.13381  
##            BrainRegion2  Treatment1:BrainRegion1  Treatment1:BrainRegion2  
##                 0.31104                 -0.03636                 -0.22824
```

```
# P2ry12
stats.input2 <- stats.input %>% filter(Antibody=="P2ry12")
stats.testing <- stats_morphologymeasures.animal(stats.input2, "Value ~ Treatment*BrainRegion", 
                                      "~Treatment|BrainRegion", "~Treatment|BrainRegion", "bonferroni")
```

```
## [1] "Foreground pixels"
## [1] "Density of foreground pixels in hull area"
## [1] "Span ratio of hull (major/minor axis)"
## [1] "Maximum span across hull"
## [1] "Area"
## [1] "Perimeter"
## [1] "Circularity"
## [1] "Width of bounding rectangle"
## [1] "Height of bounding rectangle"
## [1] "Maximum radius from hull's center of mass"
## [1] "Max/min radii from hull's center of mass"
## [1] "Relative variation (CV) in radii from hull's center of mass"
## [1] "Mean radius"
## [1] "Diameter of bounding circle"
## [1] "Maximum radius from circle's center of mass"
## [1] "Max/min radii from circle's center of mass"
## [1] "Relative variation (CV) in radii from circle's center of mass"
## [1] "Mean radius from circle's center of mass"
## [1] "# of branches"
## [1] "# of junctions"
## [1] "# of end point voxels"
## [1] "# of junction voxels"
## [1] "# of slab voxels"
## [1] "Average branch length"
## [1] "# of triple points"
## [1] "# of quadruple points"
## [1] "Maximum branch length"
## 
## Call:
## lm(formula = as.formula(paste(y.model)), data = tmp)
## 
## Coefficients:
##             (Intercept)               Treatment1             BrainRegion1  
##                17.66539                 -1.47885                  0.04940  
##            BrainRegion2  Treatment1:BrainRegion1  Treatment1:BrainRegion2  
##                 0.52513                 -0.08267                 -0.06426
```

```
stats.testing[[2]] %>% DT::datatable(., options=list(autoWidth=TRUE, scrollX=TRUE, scrollCollapse=TRUE))
```

```
do.call("grid.arrange", c(stats.testing[[4]], ncol=4)) # qqplots to check normality assumptions
```

```
stats.testing[[5]] %>% DT::datatable(., options=list(autoWidth=TRUE, scrollX=TRUE, scrollCollapse=TRUE)) # levene
```

```
stats.testing[[6]] %>% DT::datatable(., options=list(autoWidth=TRUE, scrollX=TRUE, scrollCollapse=TRUE)) # shapiro
```

```
stats.testing[[7]] # summary of model
```

```
## 
## Call:
## lm(formula = as.formula(paste(y.model)), data = tmp)
## 
## Coefficients:
##             (Intercept)               Treatment1             BrainRegion1  
##                17.66539                 -1.47885                  0.04940  
##            BrainRegion2  Treatment1:BrainRegion1  Treatment1:BrainRegion2  
##                 0.52513                 -0.08267                 -0.06426
```
