## Supplementary Info 3 for "Development of a high-throughput pipeline to characterize microglia morphological states at a single-cell resolution": SuppInfo3_Kim2023_DataAnalysis_1xLPS.html


### Kim2023\_DataAnalysis\_1xLPS

###### Jenn Kim

#### 29/10/2023

##### load in libraries

```
library(MicrogliaMorphologyR)
library(factoextra)
```

### 1XLPS (1 mg/kg) DATA ANALYSIS

##### set seed

```
set.seed(2)
```

##### load in data

```
data_1xLPS <- MicrogliaMorphologyR::data_1xLPS_mouse
```

##### exploratory data visualization and data transformation for downstream analyses

```
# gather your numerical morphology data into one column ('measure') which contains the feature name, and another column ('value') which contains measured values
data_1xLPS_gathered <- data_1xLPS %>% gather(measure, value, 7:33)

# check for outliers
outliers_boxplots(data_1xLPS_gathered)
```

```
outliers_distributions(data_1xLPS_gathered)
```

```
# checking different normalization features
normalize_logplots(data_1xLPS_gathered,1)
```

```
# transform your data in appropriate manner for downstream analyses
data_1xLPS_logtransformed <- transform_log(data_1xLPS, 1, start=7, end=33) # we will use the logtransformed data as our PCA input

# get sample size of data based on factors of interest
samplesize(data_1xLPS, Treatment, Sex)
```

```
## # A tibble: 4 × 3
## # Groups:   Treatment [2]
##   Treatment Sex     num
##   <chr>     <chr> <int>
## 1 LPS       F      4002
## 2 LPS       M      2804
## 3 PBS       F      5177
## 4 PBS       M      3310
```

##### generate heatmap of correlations across features

```
featurecorrelations(data_1xLPS, featurestart=7, featureend=33, rthresh=0.8, pthresh=0.05, title="Correlations across features")
```

#### Dimensionality reduction using PCA

```
pcadata_elbow(data_1xLPS_logtransformed, featurestart=7, featureend=33)
```

```
pca_data <- pcadata(data_1xLPS_logtransformed, featurestart=7, featureend=33,
                    pc.start=1, pc.end=10)
```

##### generate heatmap of correlations between PCs and features

You can generate a heatmap of correlations across the 27 different morphology features to investigate how they relate to each other. You can use this function to verify that features which explain similar aspects of cell morphology are more related to each other (e.g, features which describe cell area/territory span should all be highly correlated to each other compared to other features which do not).

```
pcfeaturecorrelations(pca_data, pc.start=1, pc.end=3, 
                      feature.start=17, feature.end=43, 
                      rthresh=0.75, pthresh=0.05, 
                      title="Correlation between PCs and features")
```

##### visually explore different sources of variability in dataset

```
# gather your data by experimental variables (e.g., Treatment, Sex, MouseID, etc.)
gathered_expvariables <- pca_data %>% gather(variable, value, 12:14) 

plots_expvariable(gathered_expvariables, "PC1", "PC2")
```

#### Soft clustering using Fuzzy K-means

##### prepare data for clustering

```
## for k-means clustering: scale PCs 1-3, which together describe ~85% of variability
pca_data_scale <- transform_scale(pca_data, start=1, end=3) # scale pca data as input for k-means clustering
kmeans_input <- pca_data_scale[1:3]
```

##### Cluster optimization prior to running fuzzy k-means

```
# check for optimal number of clusters using wss and silhouette methods
sampling <- kmeans_input[sample(nrow(kmeans_input), 5000),] #sample 5000 random rows for cluster optimization

fviz_nbclust(sampling, kmeans, method = 'wss', nstart=25, iter.max=50) # 4 clusters
```

```
fviz_nbclust(sampling, kmeans, method = 'silhouette', nstart=25, iter.max=50) # 4 clusters
```

#### Clustering

##### Regular k-means (hard clustering)

```
# cluster and combine with original data
data_kmeans <- kmeans(kmeans_input, centers=4)
pca_kmeans <- cbind(pca_data[1:2], data_1xLPS, as.data.frame(data_kmeans$cluster)) %>%
  rename(Cluster=`data_kmeans$cluster`)
```

##### Plot k-means clusters in PC space

```
plot <- clusterplots(pca_kmeans, "PC1", "PC2")
plot + scale_colour_manual(values=c("#BBCC33","#44BB99","#EEDD88","#EE8866"))
```

##### Cluster-specific measures on average for each morphology feature, relative to other clusters

```
clusterfeatures(pca_kmeans, featurestart=9, featureend=35)
```
